# Toward resolving gravitational effects on microbial growth with computer simulations

**DOI:** 10.64898/2026.05.15.725518

**Authors:** Andrew P. Latham, Emmanuel N. Skountzos, Stephen Lantin, Tyler Quarton, Ashwin Ravichandran, Jessica Audrey Lee, John W. Lawson

## Abstract

As the duration of space flights increases, so does the need to optimize off-planet microbial growth. Microbes can both be unintentionally brought into space and cause human disease or be intentionally harnessed for on-site bioengineering functions. However, optimizing microbial growth is challenging due to an insufficient understanding of how microbial communities are affected by the extraterrestrial environment. To address this gap, we have modified a previously developed model for cell growth in microgravity. By improving the functional form used for cell growth as well as the code usability, we enable further research into how microbial communities are influenced by gravity. Applying this model to isolate individual effects of gravity on cell growth indicates that a lack of gravity-driven flow decreases cell growth in microgravity, while the absence of sedimentation increases cell growth in microgravity. These opposite effects likely contribute to the system-dependent effects of microgravity observed experimentally.

## 1. Introduction

Human space exploration relies on carefully controlled environments to support life and ensure mission success; yet microbial communities can exhibit unpredictable behaviors within these habitats. These microbial communities are present on every mission, whether they are incidentally introduced through crew and cargo or intentionally incorporated for life-supporting functions like *in situ* resource utilization, food production, and even the astronaut gut microbiome. Spaceflight stressors, such as microgravity, elevated radiation, and ecological confinement, make predicting the behavior of these microbes even more challenging, which may impact both system stability and crew health(1). For long-duration missions, these factors raise critical concerns as microbial persistence on surfaces, enhanced biofilm formation, and increased antimicrobial resistance could compromise both crew well-being and the integrity of mission-critical systems(2; 3; 4). Further, optimization of microbe growth in space may enable the bioengineering of crucial functions like micronutrient synthesis or waste processing(5; 6; 7). While extensive research on microbial behavior in space has made progress(8; 9; 10; 11), *in situ* spaceflight studies are costly and logistically complex.

An attractive alternative to experimental space flight studies is *in silico* modeling to predict microbial behavior under various environmental conditions. While kinetic models(12), genome-scale metabolic models(13; 14; 15), integrative models(16; 17), and models based in artificial intelligence(18; 19; 20; 21; 22) have been used to describe the behavior of individual cells, agent based modeling approaches are better equipped to explore how gravity impacts microbial communities. Agent based models represent a microbial community as a group of agents whose interactions are governed by a set of rules(23; 24; 25). These rules govern how cells behave, interact with each other, and interact with their surrounding environment; consequently, modifying how cells interact with the environment provides a straightforward way to model the low gravity and/or high radiation experienced in space. Although various software packages have been developed for agent-based modeling of microbial communities(26; 27; 28; 29; 30), these models are often developed for terrestrial applications and may not be easily adaptable for exploring the effects of spaceflight. More specifically, detailed models of fluid mechanics are necessary to capture how gravity-induced fluid flow impacts cell growth.

To address this gap, **C**FD-DEM **A**rtificial **M**icrogravity **D**evelopments for **L**iving **E**cosystem **S**imulation (CAMDLES1) was introduced(31). This code extends the CFDEM®coupling modeling software(32), which couples the LAMMPS Improved for General Granular and Granular Heat Transfer Simulations (LIGGGHTS) DEM software package(33; 34) with the Open Field Operation And Manipulation (OpenFOAM) CFD software package(35). It accounts for cell growth by treating each cell as a spherical particle of variable size embedded in a fluid, represents each metabolite as a concentration field in the fluid, implements cell growth as metabolite-dependent increases to the particle radius, and describes interactions between and within the fluid and the particles using Computational Fluid Dynamics - Discrete Element Method (CFD-DEM) methods. The original study compared the growth of cross-feeding cellular colonies under microgravity, 1 g, and a rotating wall vessel, and found that cell growth in a rotating wall vessel was faster than in microgravity but slower than 1 g. However, the original study was unable to resolve previous experimental observations that observed both decreased and increased cell growth in microgravity, depending on the experimental setup(36; 37; 38; 39; 40).

Here, we explore various effects gravity may induce on cell growth using an extension of CAMDLES (CAMDLES2). The updated code improves both the underlying model for cell growth and code usability, in addition to numerous other improvements. More specifically, a Monod growth model(41) replaces the previously used Blackman growth model(42), resulting in a more realistic biological model. Additionally, the code was refactored to improve readability, and the input file format was made more flexible. Together, these advancements simplify the application of CAMDLES2 to model microbial growth that underpins key functions such as nutrient cycling, cross-feeding, and pathogenic potential, addressing a critical need in bioregenerative life support systems and risk mitigation strategies for extended space missions. We strategically applied CAMDLES2 to study different effects gravity may use to influence cell growth, and found that removing gravity-driven flow decreases cell growth, while removing sedimentation increases cell growth. These findings may partially explain the apparently contradictory results seen in previous experiments(36; 37; 38; 39; 40).

## 2. MATERIALS AND METHODS

### 2.1. CFD-DEM simulations of cell growth

CAMDLES2 is based in the CFD-DEM modeling methodology (Fig. 1). Briefly, a cell is represented as a solid, spherical particle (Table 1) whose motion is solved for by applying Newton’s laws of motion, while the fluid is represented as a grid (Table 2), whose flow is modeled by applying the Navier-Stokes equation. Values of each fluid variable are stored at the center of each CFD grid element, and values at other positions are determined through interpolation.

**Table 1:** Cell-specific variables tracked for each particle in CAMDLES2 simulations.

| Variable | Symbol [Units] |
| --- | --- |
| Cell index | $i$ |
| Cell type | $n_i$ |
| Position | $\vec{x}_i$ [$\mu\text{m}$] |
| Velocity | $\vec{v}_i$ [$\frac{\mu\text{m}}{\mu\text{s}}$] |
| Radius | $r_i$ [$\mu\text{m}$] |
| Mass | $M_i$ [pg] |
| Acetate concentration | $[A]_i$ [$\frac{\text{pg}}{\mu\text{m}^3}$] |
| Methionine concentration | $[M]_i$ [$\frac{\text{pg}}{\mu\text{m}^3}$] |

**Table 2:** Fluid-specific variables tracked on each grid center in CAMDLES2 simulations.

| Variable | Symbol [Units] |
| --- | --- |
| Kinematic pressure | $p$ [$\frac{\mu\text{m}^2}{\mu\text{s}^2}$] |
| Density | $\rho$ [$\frac{\text{pg}}{\mu\text{m}^3}$] |
| Velocity | $\vec{V}$ [$\frac{\mu\text{m}}{\mu\text{s}}$] |
| Kinematic viscosity | $\nu$ [$\frac{\mu\text{m}^2}{\mu\text{s}}$] |
| Acetate concentration | $[A]^{\text{fluid}}$ [$\frac{\text{pg}}{\mu\text{m}^3}$] |
| Methionine concentration | $[M]^{\text{fluid}}$ [$\frac{\text{pg}}{\mu\text{m}^3}$] |
| Momentum exchange between the solid and liquid phases | $K_{\text{sl}}$ [$\frac{\text{pg}}{\mu\text{m}^3 \mu\text{s}}$] |

**Figure 1:**
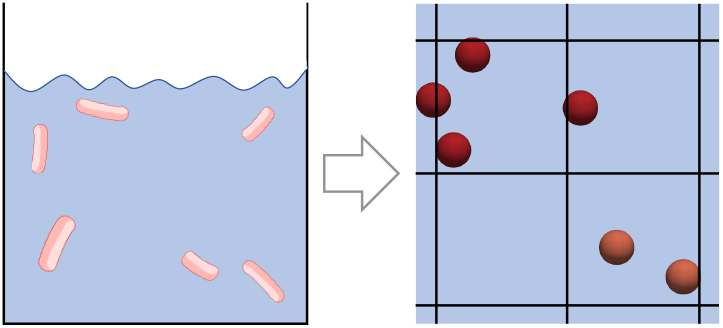
CFD-DEM model of bacterial colonies. Each cell is represented as a spherical particle, and the fluid is represented as a grid.

The DEM portion of the simulation updates the cell positions using Newton’s laws of motion. At each timestep, each cell’s velocity 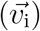 and position 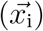 are updated in a three step velocity-Verlet algorithm(43), according to

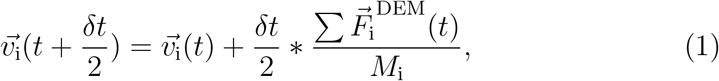

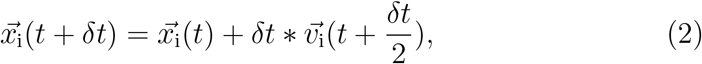

and

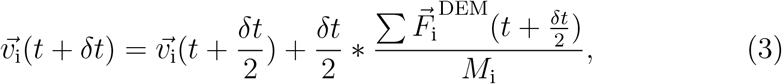

where *i* is the cell index, *M*_i_ is the cell mass, *t* is the time at the previous timestep, *δt* is the timestep, and 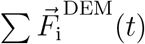 is the sum of all forces acting on the cell. In theory, these forces could include a variety of physical effects, such as interactions between cells, interactions with the fluid, and Brownian motion; although, in our simulations, we ignore the forces the fluid exerts on the cells, do not implement Brownian motion, and initialize the cells with no velocity. In practice, this holds the cells in place. This approximation was made in both this work and in CAMDLES1(31), because metabolite diffusion is several orders of magnitude (∼ 10^3^) times faster than cell diffusion(44), and our main purpose here is to explore the role of metabolite mass transport. In future work, we will explore the effects of cell dynamics.

The CFD portion of the simulations updates the flow of the fluid using the Navier-Stokes equation. We modeled the fluid as an incompressible, laminar flow and solved the system system of equations using the finite volume methods implemented in CFDEM®coupling’s “cfdemSolverPisoSTM”, a Pressure-Implicit with Splitting of Operators solver modified for coupling to DEM as well as scalar transport(45). At each timestep, the velocity of the fluid 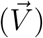 is updated by a system of linear equations of the form

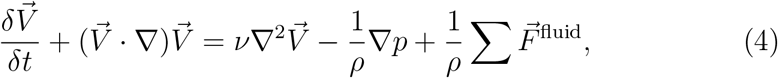

where *v* is the kinematic velocity, *ρ* is the density of the fluid, *p* is the kinematic pressure, and 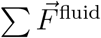 are the other forces that act on the fluid. For our system, the 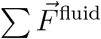 included gravity and momentum exchange between the cells and the fluid, and takes the form

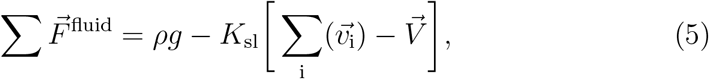

where *K*_sl_ is the momentum exchange between the solid and liquid phases, and g is the gravitational constant. These equations are solved using the type II model described by Zhou *et al* (46). These systems of equations are solved at all grid points simultaneously, allowing for an updated fluid flow at each time step.

Two way general scalar exchange accounted for transfer of metabolites between the cells and the medium. On the CFD side of the simulation, the concentration of each metabolite in the fluid (*ϕ*_f_) is updated according to a scalar transport equation of the form

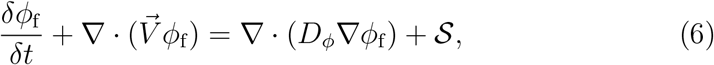

where *D*_*ϕ*_ is the diffusion coefficient of the metabolite, and 𝒮 is the source term, which determines the exchange between the fluid and the cells. This source term took the form

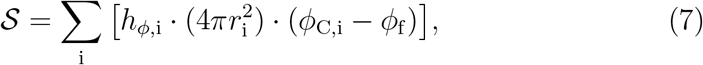

where *r*_i_ is the cell radius, *ϕ*_C,i_ is the cellular metabolite concentration and *h*_*ϕ*,i_ is the mass transfer coefficient, which is determined empirically(47), based on the volume fraction, Schmidt number, and Reynolds number. Transport into each cell is proportional to the local metabolite concentration difference between the fluid and the cell, taking the form

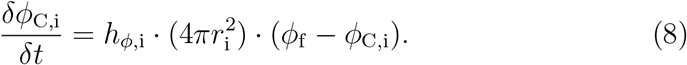

### 2.2. Monod growth model

In CAMDLES1, a Blackman growth model was used to approximate the growth of each cell(42). This growth model increased the cell mass by a constant rate, as long as sufficient substrate concentration was present. When the concentration was below the cutoff substrate concentration, the cell grew proportional to the amount of substrate present.

To make the cell growth model more realistic, CAMDLES2 implements a Monod growth model for each cell, which is frequently used in metabolic cell modeling(41). Although both Blackman and Monod capture substrate dependent growth at low substrate concentration and substrate independent growth at high substrate concentration, Monod is often more accurate, likely due to the sharp transition seen in Blackman growth(48). Compared to Blackman growth, Monod growth predicts faster growth at low substrate concentrations, and slower growth at high substrate concentrations (Fig. 2). The updating equations for the Monod model are implemented at each cell and take the form:

**Figure 2:**
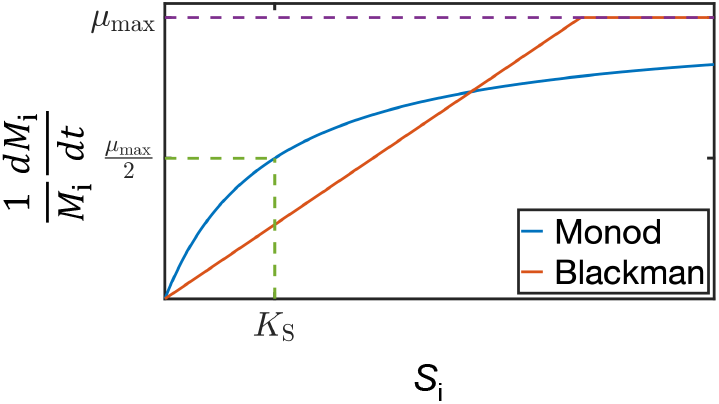
Example of growth models. The solid lines show cell growth rates of Monod and Blackman growth models, while dashed lines highlight the parameters of the Monod model (*K*_S_, *µ*_max_).

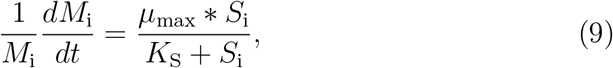

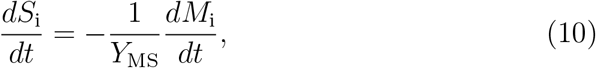

and

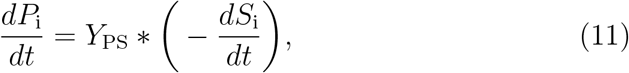

where *M*_i_ is the mass of the cell, *S*_i_ is the cellular substrate (consumed metabolite) concentration, *P*_i_ is the cellular product (secreted metabolite) concentration, *µ*_max_ is the maximum growth rate of the cell, *K*_S_ is the substrate concentration at which the growth rate is half of *µ*_max_, *Y*_MS_ is the growth yield coefficient, which represents the amount of *M*_i_ produced per *S*_i_ consumed, and *Y*_PS_ is the yield of *P*_i_ produced per *S*_i_ consumed. Here, *µ*_max_, *K*_S_, *Y*_MS_, and *Y*_PS_ are assumed to be constants. The radius of the cell is calculated based on the change in cell mass by assuming a constant mass density 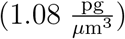 as the cell grows. Note that *S*_i_ and *P*_i_ are cellular metabolite concentrations, and thus are instances of *ϕ*_C,i_ (Eq. 7-8), which describes how these variables are coupled to their respective fluid concentrations.

In order to accelerate the timescales of our simulations, cells start from the same size and grow in radius but do not divide. For the same reason, modeled growth rates were larger than biological growth rates to observe mass transport-limited effects within our simulation timescales.

### 2.3. System details

While many biological systems could be modeled with CAMDLES2, we chose to focus on a simple system of cross-feeding bacteria. Specifically, a community of *Escherichia coli* (*E. coli*) and *Salmonella enterica* (*S. enterica*) that have been genetically engineered for cooperation(49; 50). The modified *E. coli* are unable to produce methionine, and depend on methionine produced by the *S. enterica* for growth. In exchange, when grown on lactose, the *E. coli* produce a carbon byproduct needed for growth by the *S. enterica*, which cannot metabolize lactose. Here, we use a simplified version of this system in which *E. coli* consume methionine and secrete acetate while *S. enterica* consume acetate and secrete methionine as a test case to examine how reduced flow from a lack of gravity effects the cross-feeding behavior and, thus, cell growth. We use the term “metabolite” to refer to methionine and acetate collectively.

For this work, simulations tested the cell growth dependence on initial metabolite concentration, gravity, and cell configuration. Five initial metabolite concentrations 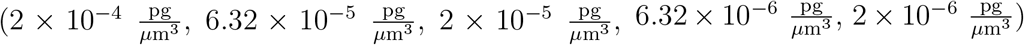, two gravitational conditions (no gravity and terrestrial gravity), and three starting configurations (uniform, 2-colony, and sedimented) were used. For metabolites, the starting concentrations of acetate and methionine were identical to each other in each simulation, and metabolites were uniformly distributed at the specified concentration both throughout the fluid and within all cells. The lowest metabolite concentration 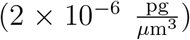 was used unless otherwise noted and was chosen such that the concentration was similar to *K*_S_, which would start at a growth rate of approximately 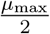. Meanwhile, the other concentrations were chosen to investigate the system’s behavior as the growth rate approached ∼*µ*_max_ (Fig. 2). Simulations with gravity used a gravitational constant of 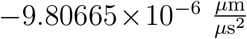, while no gravity simulations used a gravitational constant of 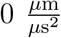. All simulations were conducted in a square simulation box with sides lengths of 300 *µ*m. Simulations with uniform starting configurations had cells uniformly distributed throughout the entire simulation box; 2-colony simulations had *E. coli* and *S. enterica* cells uniformly distributed within separate 100*µ*m × 100*µ*m × 100*µ*m boxes, whose center was positioned 259.81*µ*m away from each other; and sedimented simulations had cells uniformly distributed throughout the entire X and Y dimension and in the lower quarter of the box in the Z dimension. Periodic boundaries were used for uniform and 2-colony configurations, while sedimented configurations and their comparable simulations with uniform configurations used fixed boundaries, which prevents fluid exchange between opposite ends of the simulation box. Each cell configuration results in a different local cell concentration, despite the overall cell concentration being maintained.

Other parameters used in this work were constant across all simulations. Simulations included 2000 cells, 1333 of which were *E. coli* cells and 667 of which were *S. enterica* cells to mimic the steady state species ratio observed empirically in our lab. All cells were held stationary during the simulations, meaning they did not move within the simulation box and forces the fluid exerted on the cells were ignored, as it was done in CAMDLES1(31), because metabolite diffusion is several orders of magnitude (∼10^3^) times faster than cell diffusion(44). Despite cells not moving, they still exert forces on the fluid. Detailed parameters for Monod cell growth (Table S1), the CFD portions of the simulation (Table S2), the DEM portion of the simulation (Table S3), and the coupling between the CFD and DEM portions of the simulation (Table S4) can be found in the Supplementary Information.

### 2.4. Benchmarking

To test our computational model, we performed two simple tests to ensure the modifications we made to CFDEM®coupling produce the expected results. In a first test, we checked the gravity model. We turned off all forces acting on the fluid except gravity, and performed three short simulations at arbitrary values of the gravitational constant. In each case, the fluid velocity matches the analytical expectation (Fig. S1), indicating that the gravity functions as expected. In a second test, we examined cell growth, metabolite consumption, and metabolite production. We created a simple system with one type of each cell, placed far apart in a simulation box. In this setup, we performed a short simulation. Each quantity matched the analytical form of the Monod equation (Eqs. 9-11), indicating the cell growth code performs as expected (Fig. S2).

After benchmarking our setup in simple cases, we wanted to observe the change in system mass in simulations similar to those used throughout the manuscript. As we set *Y*_MS_ +*Y*_PS_ = 1 (Eqs. 9-11), we expected the total mass of the system to be conserved during the simulation. In realistic systems, the total yield for the metabolites considered here may actually be less than unity due to the production of other species; however, we make this choice here for simplicity. To test this hypothesis, we tracked the mass of metabolites and cells in a simulation with uniform cell position, low metabolite concentration, gravity, and periodic boundaries (Fig. S3). Our results suggest that the total mass of the system remains nearly-constant, and that metabolites are converted into cell mass throughout the simulation. Methionine is consumed more quickly than acetate, because our system has twice as many *E. coli* cells (which consume methionine) as *S. enterica* cells (which consume acetate).

### 2.5. Code

CAMDLES2 improves on CAMDLES1 by making the code easier to use for other researchers, in addition to other improvements. The source code is included as text in this manuscript’s Supplementary Information (*Source Code*). A guide for code installation is provided in the Supplementary Information (*Installation Guide*). Simulations are run by calling source `Allrun.sh > display`. An example system setup with no gravity, low starting metabolite concentration, and periodic boundary conditions is included as text in the Supplementary Information, along with instructions with how to derive the other systems from this example (*Example*). The code to write initial configurations and the code to analyze the radius of the cell and the cellular concentration of metabolites are also included as text in the Supplementary Information (*Analysis and Initial Configurations*). Paraview was used to visualize the simulations(51).

## 3. RESULTS AND DISCUSSION

We conducted a series of simulations using CAMDLES2 to examine the effect of gravity on bacterial growth in a cross-feeding microbial community, starting by looking at interactions between gravity and other environmental conditions such as initial metabolite concentration and the spatial distribution of cells. These simulations focus on the growth of cross-feeding bacteria, *E. coli* and *S. enterica*(49; 50). To best isolate how each variable contributes to cell growth, the main text focuses on comparisons to a system with low starting metabolite concentration, no gravity, and uniform starting configuration unless otherwise noted. We focus on results in terms of average *E. coli* radius and average methionine concentration in *E. coli* cells, as average *S. enterica* radius and average acetate concentration in *S. enterica* cells obey identical trends in the parameter regimes tested. As differences in values between different gravitational conditions or starting configurations were quite small, we also compared ratios of these quantities.

### 3.1. Gravity accelerates bacterial growth

To begin, we focused on how gravity influences the cell growth. As was done in CAMDLES1(31), we intentionally remove gravitational effects from acting on the cells. As a result, cells are held in place; meanwhile, gravity acts on the fluid. In this way, our model can uniquely isolate effects from either mass transport in the fluid or cell configuration. In simulations with gravity, the velocity of the fluid increases over the course of the simulation, but the velocity remains at 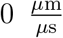 in simulations without gravity (Fig. 3a). The velocity in the gravity simulation is near-uniform, as the only heterogeneity arises due to friction from individual cells.

**Figure 3:**
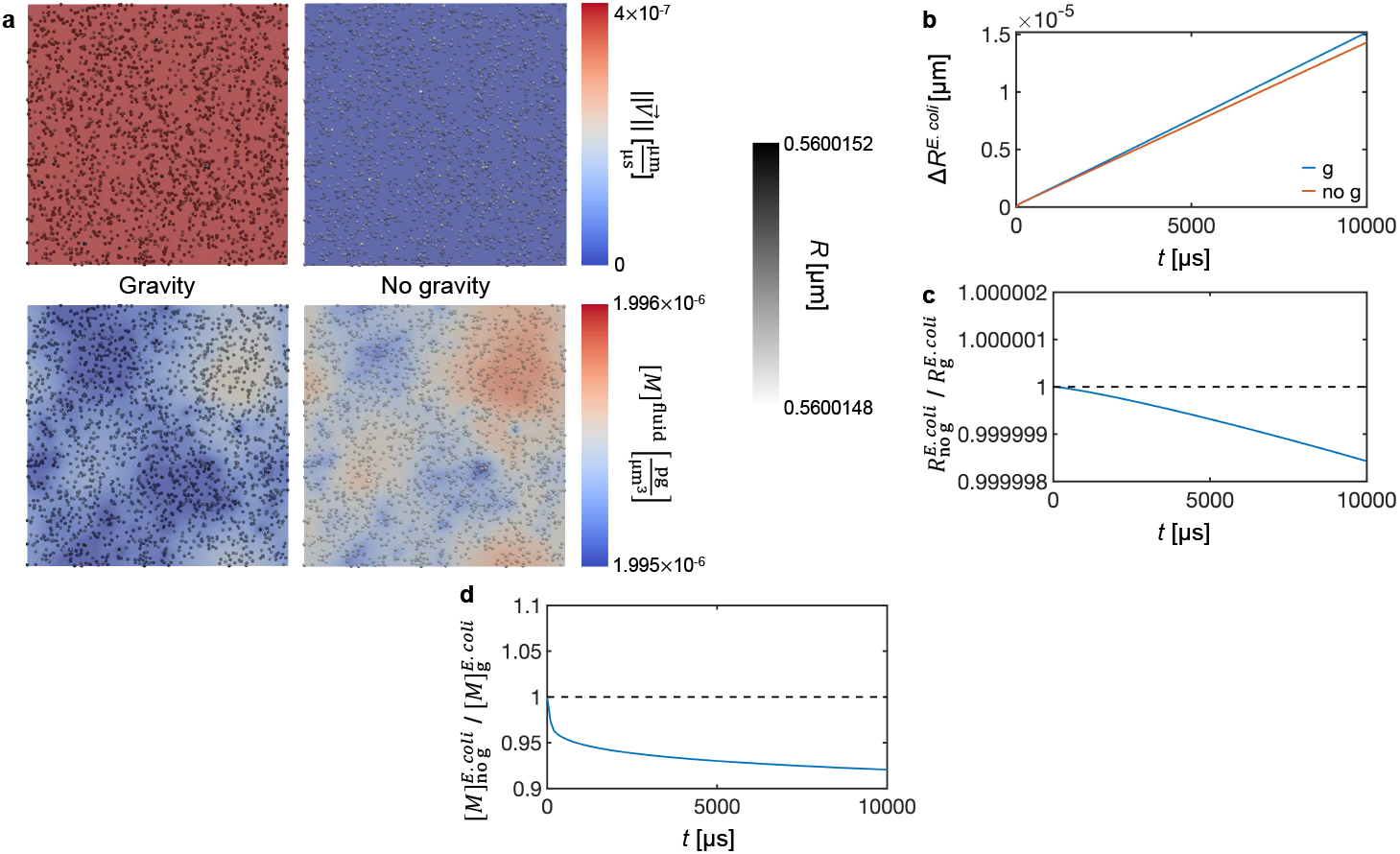
Comparison between simulations in gravity and no gravity conditions, at low metabolite concentration and uniform cell distribution. (a) Images of simulations with (left) and without (right) gravity. The solution is colored by either the magnitude of the fluid velocity (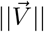, top) or the fluid methionine concentration ([*M*]^fluid^, bottom), while the cells are colored by the radius (R). The average change in *E. coli* radius (Δ*R*^*E.coli*^) (b), the ratio between the average *E. coli* radius without gravity 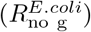 and with gravity 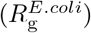 (c), and the ratio between the average methionine concentration in *E. coli* without gravity 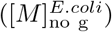 and with gravity 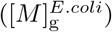 (d) as a function of time. Shaded region in (b) represent the standard deviation over all cells and is on the scale of the linewidth when not visible.

The results suggest that cell growth decreases in no gravity conditions relative to when gravity is present (Fig. 3b-c). This result arises due to lower cellular metabolite concentration (Fig. 3c, S4a), which is caused by a lack of fluid mixing in no g. In both gravity and no gravity conditions, the cellular metabolite concentration starts at 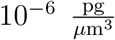 creases to 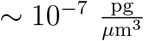 due to metabolite consumption for cell growth. After this initial sharp decrease, the concentration remains at a near steady state that balances metabolite consumption during cell growth and intake from the fluid. As metabolites circulate through the fluid more slowly without gravity than with gravity, cells in no gravity conditions replenish their metabolite concentration more slowly, leading to a lower cellular metabolite concentration. The decreased growth rate leads to decreased metabolite consumption, resulting in a higher methionine concentration in the fluid without gravity than with gravity (Fig. S4b). These findings do not depend on the length of the simulation (Fig. S5).

To test the effect of initial metabolite concentration on cell growth, we conducted simulations at five different metabolite concentrations, starting at ∼ *K*_S_ (2 × 10^*−*6^ *µ*s^*−*1^), with each subsequent concentration increasing by half an order of magnitude in log-space (Fig. 4a). Interestingly, we observed a non-monotonic effect on the difference in growth rate between gravity and no gravity conditions with increasing initial concentration (Fig. 4b). At smaller concentrations 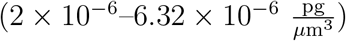, the difference in growth rate increases with increasing initial metabolite concentration. This trend seems to be caused by increased differences in metabolite intake, for the metabolite concentration ratios follow the same pattern (Fig. 4c). These results can be rationalized by the formulas in our model: both cell growth (Eq. 9) and metabolite intake (Eq. 8) increase with increasing cell radius; therefore, as cells grow more quickly at increasing metabolite concentration, differences in growth accumulate more quickly as well. However, this trend is reversed as the initial concentration continues to increase 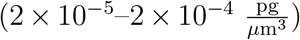. This reversal is likely caused by cell growth approaching its maximum growth rate, *µ*_max_ (Eq. 9). At low initial concentrations, the *E. coli* methionine concentration drops rapidly, until a steady-state is reached that balances consumption for cell growth and intake from the solution (Fig. S6a). At high initial metabolite concentrations, the drop in the *E. coli* methionine is negligible, meaning cell growth can maintain a near maximum rate (Fig. S6b). As a near maximum growth rate can be maintained with or without gravity, the effect of gravity becomes smaller. This decrease of the effect of gravity at high metabolite concentrations has previously been seen experimentally, where the gravity-dependence of *Pseudomonas aeruginosa* growth disappeared at high phosphate availability(40). Together, these findings suggest metabolite concentration could be engineered to control microgravity’s impact on cell growth.

**Figure 4:**
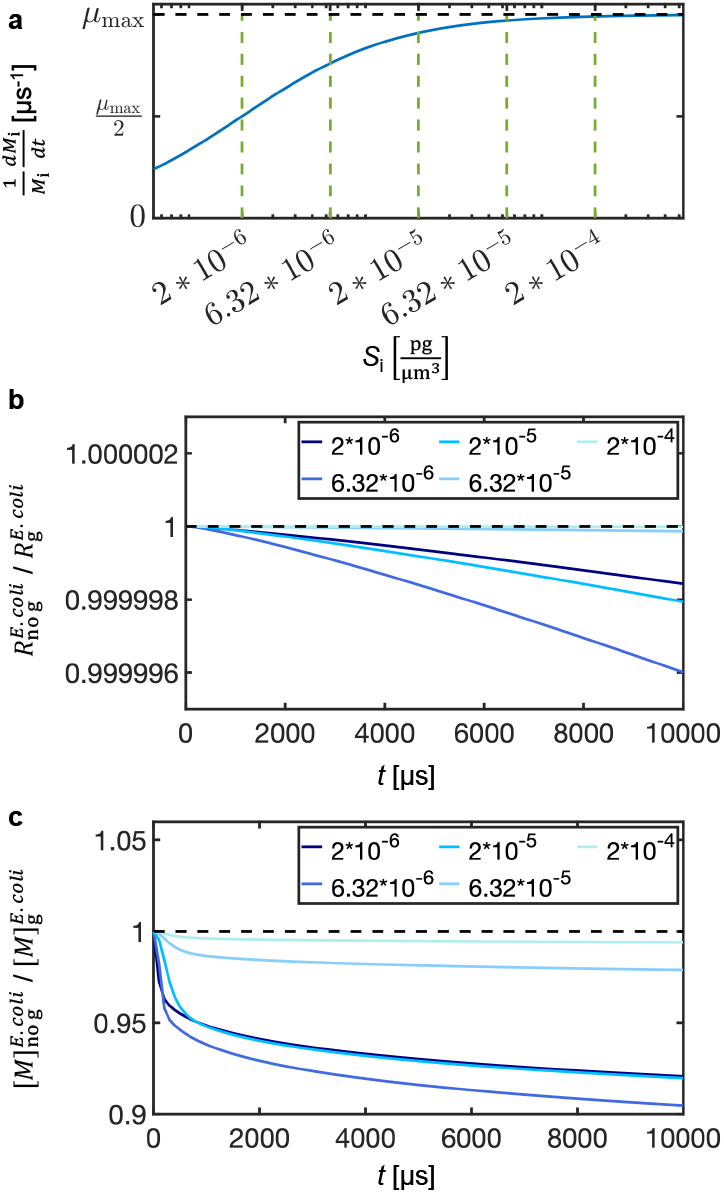
How varying the initial metabolite concentration influences the effect of gravity on cell growth, with uniform cell distribution. (a) Monod growth rate plotted as a function of substrate concentration, with the parameters used in this model. Vertical dashed lines highlight the starting concentrations used in these simulations, while the horizontal dashed line denotes µ_max_. (b) The ratio between the average *E. coli* radius without gravity 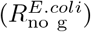 and with gravity 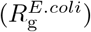, and (c) the ratio between the average methionine concentration in *E. coli* without gravity 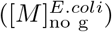 and with gravity 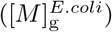. Initial metabolite concentrations are indicated in the legends 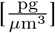.

### 3.2. Separating cells into colonies slows bacterial growth

After examining the effects of gravity on cell growth, we focused on the effect of starting position. In these simulations, cells were placed either uniformly throughout the simulation box (uniform) or in two distinct colonies, each of which has only one cell type (2-colony), and simulated in no gravity conditions (Fig. 5a). The 2-colony simulation created a concentration gradient in which methionine appears at lower concentration than the bulk near the *E. coli* colony and at higher concentration than the bulk near the *S. enterica* colony. This concentration gradient emerges from the properties of the system, as methionine is produced by *S. enterica* and consumed by *E. coli*. Thus, the methionine must diffuse away from the *S. enterica* colony before it can be consumed by the *E. coli* colony. Furthermore, this gradient is reversed for acetate, which is produced by *E. coli* and consumed by *S. enterica* (Fig. S7).

**Figure 5:**
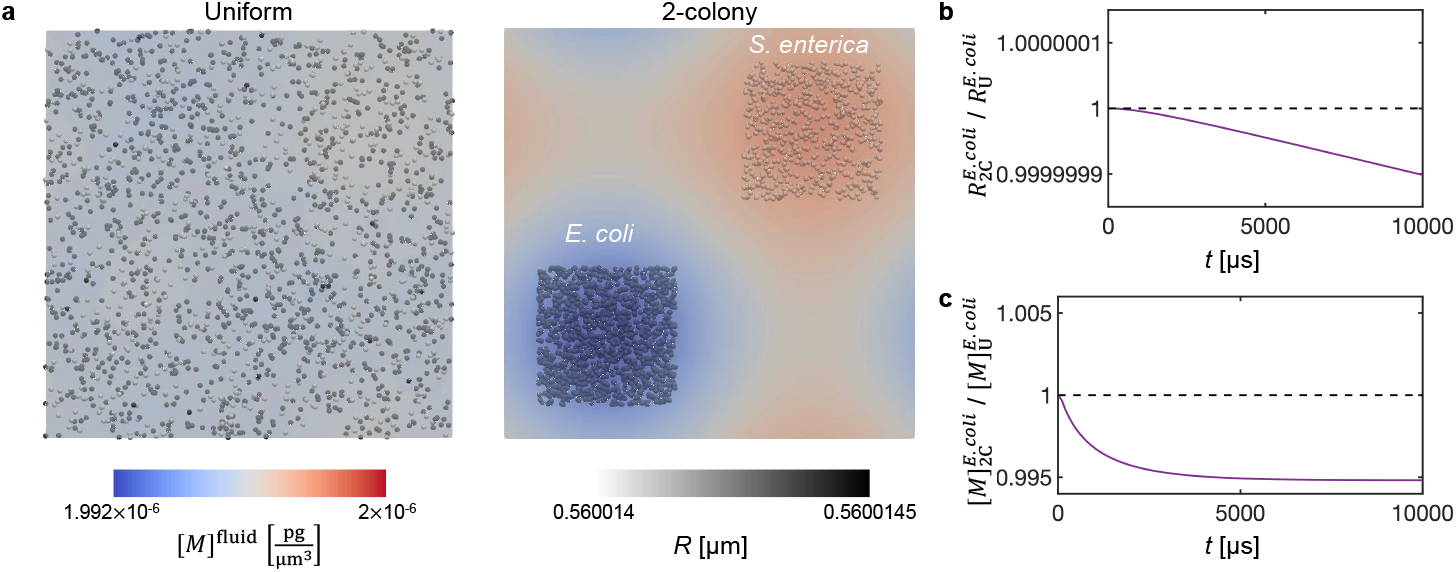
Comparison between uniform and 2-colony simulations at low metabolite concentration and no gravity. (a) Images of uniform (left) and 2-colony (right). The solution is colored by the methionine concentration, while the cells are colored by the radius. The ratio between the average *E. coli* radius in 2-colony 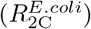 and uniform 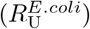 start-ing positions (b), and the ratio between the average methionine concentration in *E. coli* from 2-colony 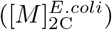 and uniform 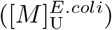 starting positions (c) as a function of time.

As a result of the two-step process of cross-feeding, where production is followed by diffusion, 2-colony grows slower (Fig. 5b) and has a lower methionine concentration in the *E. coli* colony (Fig. 5c) and a lower acetate concentration in the *S. enterica* colony than uniform.

### 3.3. Sedimentation rationalizes differing experimental observations on how removing gravity perturbs bacterial growth

Thus far, we have shown that removing gravity decreases cell growth due to the lack of gravity-driven flow. However, both increased and reduced growth have been observed in experimental studies of microbe growth under microgravity(10). This apparent contradiction has been partially attributed to gravity-induced sedimentation as a counteracting force that would increase cell growth without gravity, in addition to other mechanisms such as changes to the cell cycle(11). Therefore, we used CAMDLES2 to test the effect of gravity-induced sedimentation on cell growth.

To create an environment capable of studying sedimentation, we removed periodic boundary conditions (PBC, Fig. S8). PBC allow cells or fluid that exits on one side of the simulation box to reenter on the other side, creating an effectively infinite system. With PBC and with gravity, the velocity of the system initially increases in a spatially uniform manner, while the pressure is negligible. However, replacing periodic boundaries with fixed boundary conditions (FBC) inherently creates velocity and pressure gradients in the simulation box. The walls slow the flow of the fluid, so that the velocity is larger near the middle of the simulation box than near the edges. Further, the motion of the fluid creates a pressure gradient, with negative pressure at the top of the simulation box and no pressure near the bottom. Although not accessible on our simulation timescale, the long-time behavior of these systems would also differ: the net velocity of a fluid in PBC would converge to a terminal velocity, while the net velocity of a fluid in FBC would converge to zero.

To mimic the effect of sedimentation, the same number of cells from the uniform case were placed in the bottom quarter of the box, effectively increasing the local density of the cells. In this setup, simulations were performed without gravity and with FBC and compared to simulations in uniform cell positions without gravity and with FBC. The sedimented system creates a metabolite gradient, as metabolite consumption creates a metabolite deficient region around the cells, which is seen in both methionine (Fig. 6a) and acetate (Fig. S9). This symmetric gradient is different than the opposing gradients seen in the two colony case, because both cell types are now mixed. Thus, growth in sedimented cells is slower than for uniformly distributed cells (Fig. 6b), due to increased metabolite availability in the uniform case (Fig. 6c). The lower metabolite availability in sedimented cells is due to the metabolite gradient; the local metabolite concentration near the sedimented cells is replenished by production from the other cell type and diffusion from the cell-free region of the simulation box, which effectively serves as a metabolite reservoir. This is also evident in analysis of the fluid metabolite concentration, which shows the average methionine concentration in the fluid is nearly identical in uniform and sedimented conditions, but the variance of the methionine concentration across the fluid is much larger in sedimented than uniform (Fig. 6d), due to larger concentrations than the mean far from the sedimented cells and smaller concentrations than the mean close to the sedimented cells. These conclusions are not changed when gravity is included in the system (Fig. S10). Thus, cells grow faster before they are sedimented than they do after they are sedimented, due to a high local ratio of cells to metabolites. This effect may serve as a mechanism through which microgravity would increase the rate of cell growth; although, this observation does not account for the growth rate during the sedimentation process.

**Figure 6:**
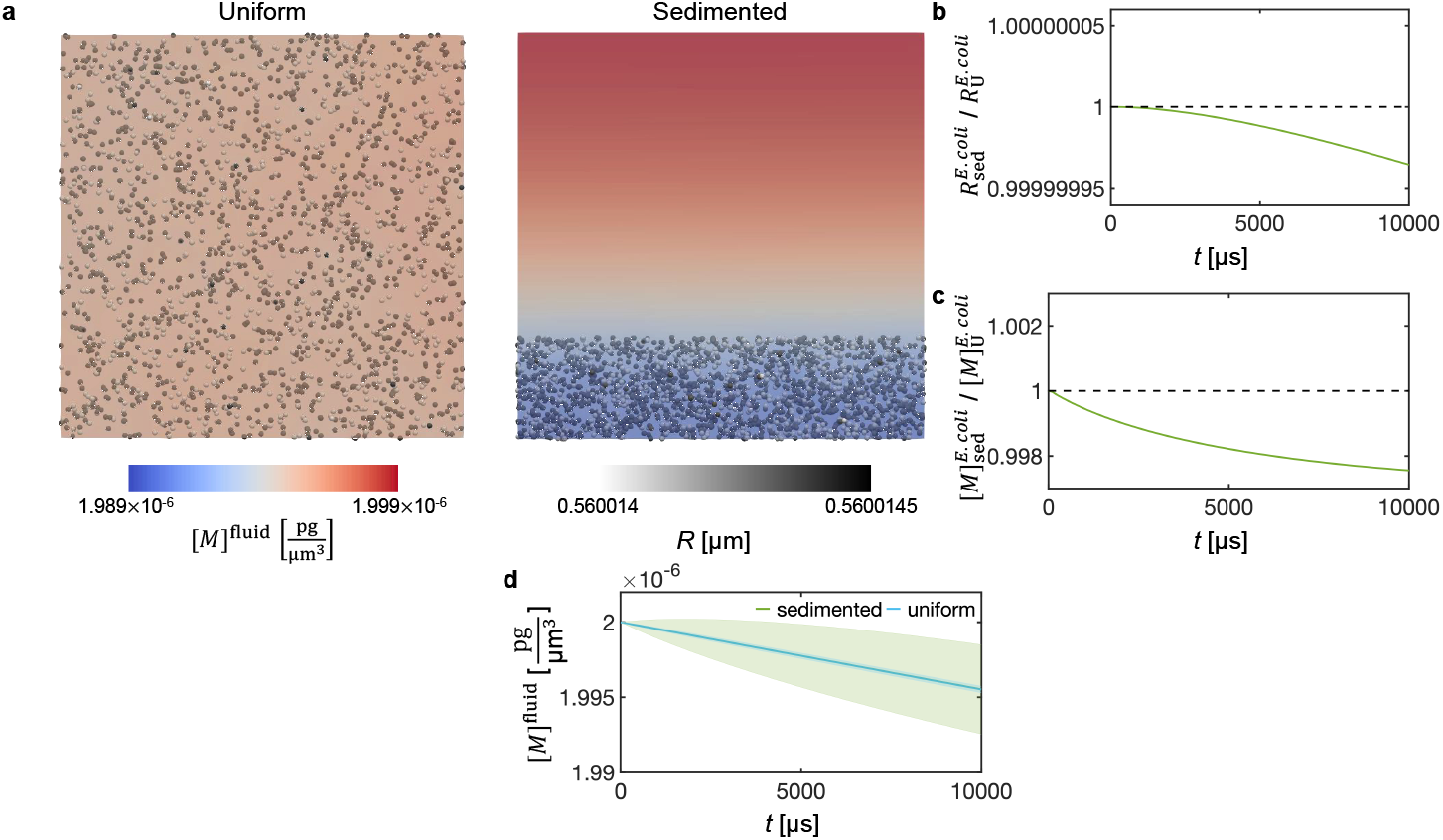
Comparison between simulations starting from uniform or sedimented starting positions with low metabolite concentration, no gravity and FBC. (a) Images of simulations with uniform (left) and sedimented (right) positions. The solution is colored by the methionine concentration, while the cells are colored by the radius. The ratio between the average *E. coli* radius in sedimented 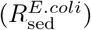 and uniform 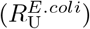 starting positions (b), the ratio between the average methionine concentration in *E. coli* from sedimented 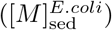 and uniform 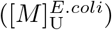 starting positions (c), and the average methionine concentration in the fluid (d) as a function of time. Shaded region in (d) represent the standard deviation over all grid positions.

## 4. CONCLUSIONS

The present work presents a new version of the CAMDLES simulation code, a CFD-DEM tool for studying microbial communities. This new version, CAMDLES2, improves the analytical form for cell growth while also enhancing the usability and readability of the code. To demonstrate the power of CAMDLES2, we applied it to isolate the physical mechanisms through which gravity may perturb cell growth. Removing gravity-induced flow decreased cell growth rate; moreover, the magnitude of this effect could be controlled by the metabolite concentration. This is in agreement with the long-held hypothesis that the lack of convection can result in stress responses in cells cultured in microgravity(11). Meanwhile, our model predicts that a lack of gravity-induced sedimentation would increase cell growth. The balance of these reverse effects likely explains the difficulty in determining simple rules for how cell growth is effected by microgravity(10).

In the future, we hope to continue to develop CAMDLES and apply it to study biologically interesting systems. In particular, the biological side of the model can be improved by incorporating known biophysical features such as a more complex cell metabolism, cell motility, cell division, and chemotaxis. Further, the physics of the model could be enhanced through implementing multiple versions of a rotating wall vessel, mirroring different experimental setups. Such implementations would enable direct comparison between simulation and experiment, advancing both experimental design and our understanding of biology under microgravity.

## 5. GOVERNMENT RIGHTS NOTICE

This work was authored by employees of KBR Wyle Services, LLC under contract no. 80ARC020D0010 with the National Aeronautics and Space Administration. The United States Government retains and the publisher, by accepting the article for publication, acknowledges that the United States Government retains a nonexclusive, paid-up, irrevocable, worldwide license to reproduce and prepare derivative works, distribute copies to the public, perform publicly, and display publicly or allow others to do so for United States Government purposes. All other rights are reserved by the copyright owner. The authors declare no competing financial interest.

## 6. ACKNOWLEDGEMENTS

We acknowledge funding support from the NASA Ames Research Center - Center Innovation Fund (CIF). S.L. was further supported by the NASA Space Technology Graduate Research Opportunities (NSTGRO) Program Grant #80NSSC21K1257. The computational resources provided by NASA Advanced Supercomputing Center (NAS) is gratefully acknowledged.

