## Supplementary Information for "Toward resolving gravitational effects on microbial growth with computer simulations"

#### Contents

**Supplemental Figures** **S4**

Figure S1 . . . . . . . . . . . . . . . . . . . . . . . . . . . . . . . . . . . . S4

Figure S2 . . . . . . . . . . . . . . . . . . . . . . . . . . . . . . . . . . . . S5

Figure S3 . . . . . . . . . . . . . . . . . . . . . . . . . . . . . . . . . . . . S6

Figure S4 . . . . . . . . . . . . . . . . . . . . . . . . . . . . . . . . . . . . S7

Figure S5 . . . . . . . . . . . . . . . . . . . . . . . . . . . . . . . . . . . S8
Figure S6 . . . . . . . . . . . . . . . . . . . . . . . . . . . . . . . . . . . S9
Figure S7 . . . . . . . . . . . . . . . . . . . . . . . . . . . . . . . . . . . S10
Figure S8 . . . . . . . . . . . . . . . . . . . . . . . . . . . . . . . . . . . S11
Figure S9 . . . . . . . . . . . . . . . . . . . . . . . . . . . . . . . . . . . S12
Figure S10 . . . . . . . . . . . . . . . . . . . . . . . . . . . . . . . . . . S13

**Supplemental Tables** **S14**

Table S1 . . . . . . . . . . . . . . . . . . . . . . . . . . . . . . . . . . . S14
Table S2 . . . . . . . . . . . . . . . . . . . . . . . . . . . . . . . . . . . S15
Table S3 . . . . . . . . . . . . . . . . . . . . . . . . . . . . . . . . . . . S16
Table S4 . . . . . . . . . . . . . . . . . . . . . . . . . . . . . . . . . . . S17

**Installation Guide** **S18**

Ubuntu 18.04 . . . . . . . . . . . . . . . . . . . . . . . . . . . . . . . . . S18
Red Hat Enterprise Linux (RHEL) . . . . . . . . . . . . . . . . . . . . . . . S21
Arch Linux - EndeavourOS . . . . . . . . . . . . . . . . . . . . . . . . . . S26
Linux virtualization on macOS and Windows . . . . . . . . . . . . . . . . . . S27

**Supplemental Code** **S28**

Source Code . . . . . . . . . . . . . . . . . . . . . . . . . . . . . . . . . S28
fix_cfd_coupling_convection_species.cpp . . . . . . . . . . . . . . . . . . S30
fix_cfd_coupling_convection_species.h . . . . . . . . . . . . . . . . . . . S44
fix_scalar_transport_equation.cpp . . . . . . . . . . . . . . . . . . . . . . S48
fix_scalar_transport_equation.h . . . . . . . . . . . . . . . . . . . . . . . S77
cfdemSolverPisoSTM.C . . . . . . . . . . . . . . . . . . . . . . . . . . . . S83
Analysis and Initial Configurations . . . . . . . . . . . . . . . . . . . . . S94
analyze_DEM.py . . . . . . . . . . . . . . . . . . . . . . . . . . . . . . . . S96
write_data_uniform.py . . . . . . . . . . . . . . . . . . . . . . . . . . . . S113

write_data_2colony.py . . . . . . . . . . . . . . . . . . . . . . . . . . . . . S117
write_data_sedimentation.py . . . . . . . . . . . . . . . . . . . . . . . . . . S122
Example . . . . . . . . . . . . . . . . . . . . . . . . . . . . . . . . . . . . . . . . . S126

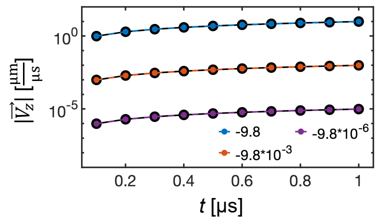


Figure S1: Testing the model for gravity. The magnitude of the fluid's velocity Z-vector ($|\vec{V_z}|$) is plotted as a function of time. Plots are made using three different strengths of the gravitational acceleration, as labeled in the legend [$\frac{\mu m}{\mu s^2}$], and are compared to the analytical expression (black lines). These simulations use a version of the model where gravity is the only force exerted on the fluid.

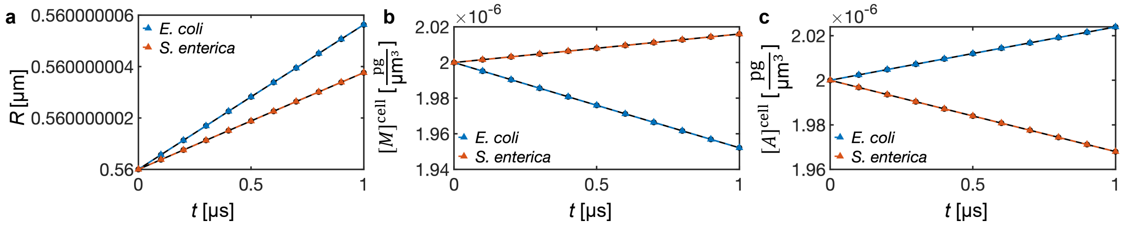


Figure S2: Testing cell growth, metabolite consumption, and metabolite production. The radius of each cell ($R$, a), the concentration of methionine in each cell ($[M]^{\text{cell}}$, b), and the concentration of acetate in each cell ($[A]^{\text{cell}}$, c) is compared to the analytical result from the Monod equation (Eq. 9-11, black line). These simulations have one *E. coli* cell at [100,100,100] $\mu$m and one *S. enterica* cell at [200,200,200] $\mu$m in a square simulation box with sides lengths of 300 $\mu$m. Cells start a with metabolite concentrations of $2 \times 10^{-6}$ $\frac{\text{pg}}{\mu\text{m}^3}$ for each metabolite, no metabolites are in the fluid, and the metabolite diffusion coefficients are set to $10^{-20}$ $\frac{\mu\text{m}^2}{\mu\text{s}}$. These changes were made to isolate the effects of the Monod equation from the fluid mechanics, and other parameters are identical to those used throughout the manuscript.

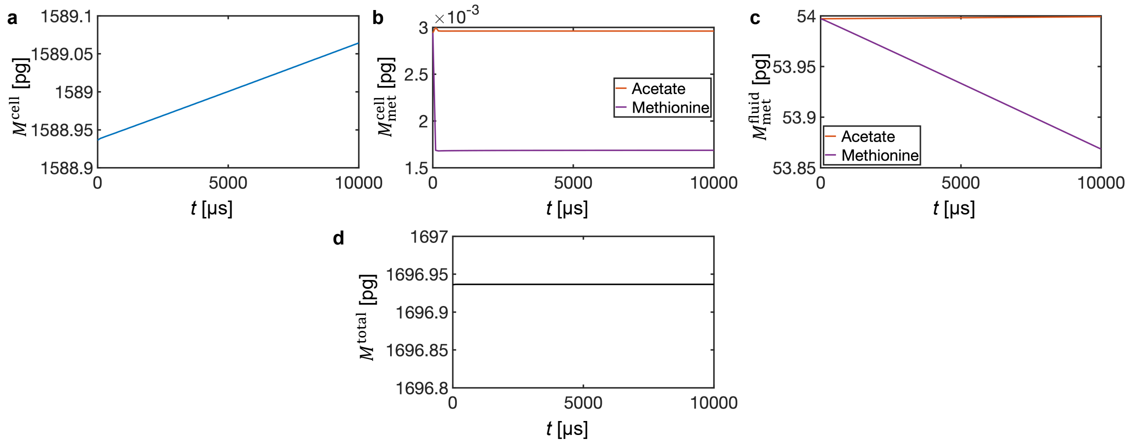


Figure S3: Testing mass conservation in a simulation with uniform cell position, low metabolite concentration, gravity, and periodic boundaries. The total mass of *E. coli* and *S. enterica* cells ($M^{\mathrm{cell}}$, a), the mass of metabolites in the cell ($M^{\mathrm{cell}}_{\mathrm{met}}$, b), the mass of metabolites in the fluid ($M^{\mathrm{fluid}}_{\mathrm{met}}$, c), and the total mass of the system ($M^{\mathrm{total}}$, d) are plotted as a function of time.

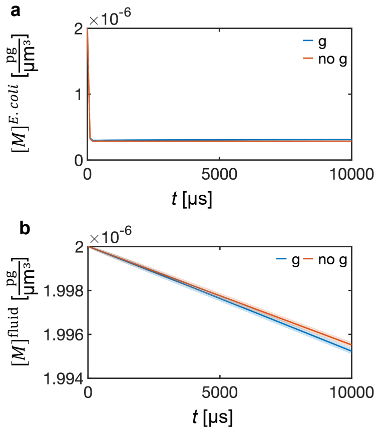


Figure S4: Additional comparison between simulations in gravity and no gravity conditions, at low metabolite concentration and uniform cell distribution. The average cellular methionine concentration in *E. coli* ($[M]^{E.coli}$) (a), and the average methionine concentration in the fluid ($[M]^{\text{fluid}}$) (b) as a function of time. Shaded regions in (a) and (b) represent the standard deviation over all cells or grid positions, and is on the scale of the linewidth when not visible.

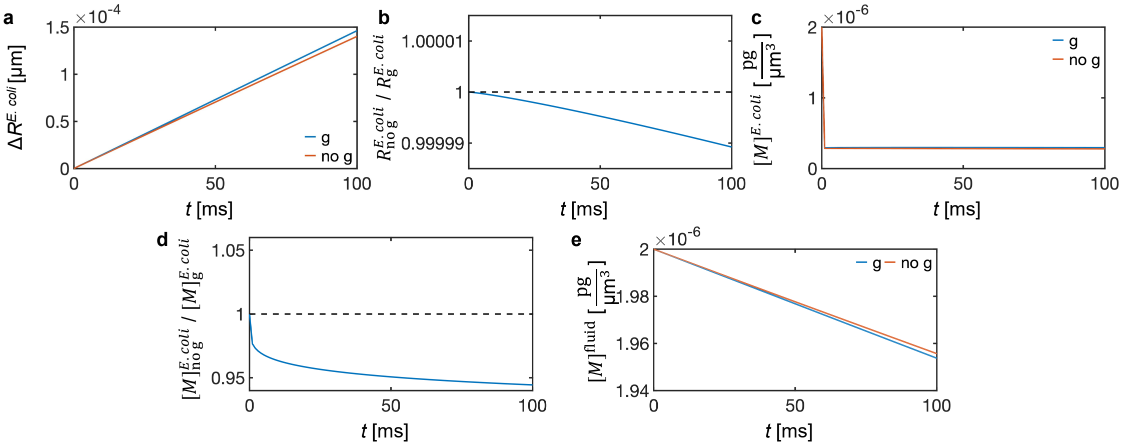


Figure S5: Extending simulation length does not change the effect of gravity. Simulations are performed from uniform configurations with and without gravity at low metabolite concentration and uniform cell distribution for ten-times longer than other simulations. The average change in *E. coli* radius ($\Delta R^{E.coli}$) (a), the ratio between the average *E. coli* radius without gravity ($R^{E.coli}_{\text{no g}}$) and with gravity ($R^{E.coli}_{\text{g}}$) (b), average cellular methionine concentration in *E. coli* ($[M]^{E.coli}$) (c), the ratio between the average methionine concentration in *E. coli* without gravity ($[M]^{E.coli}_{\text{no g}}$) and with gravity ($[M]^{E.coli}_{\text{g}}$) (d), and the average methionine concentration in the fluid ($[M]^{\text{fluid}}$) (e) as a function of time. Shaded region in (a), (c), and (e) represent the standard deviation over all cells or grid positions, and is on the scale of the linewidth when not visible. These simulations use a CFD bin size of $10\mu\text{m} \times 10\mu\text{m} \times 10\mu\text{m}$ to improve computational efficiency.

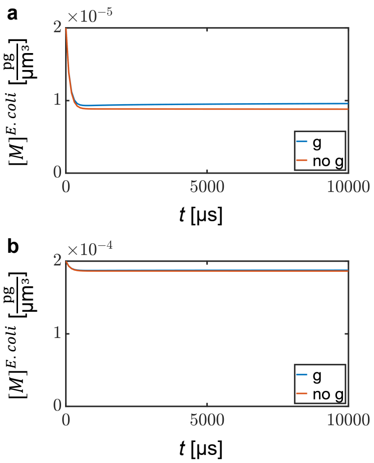


Figure S6: Additional data on how varying the initial metabolite concentration influences the effect of gravity on cell growth, with uniform cell distribution. The average cellular methionine concentration in *E. coli* ($[M]^{E.coli}$) with initial concentrations of $2 \times 10^{-5} \frac{\text{pg}}{\mu\text{m}^3}$ (a) and $2 \times 10^{-4} \frac{\text{pg}}{\mu\text{m}^3}$ (b).

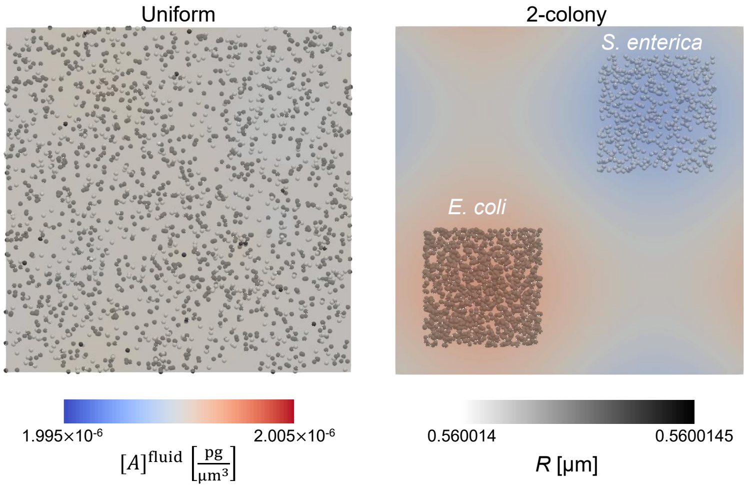


Figure S7: Comparison between uniform and 2-colony simulations. Images of uniform (left) and 2-colony (right) simulations with the solution colored by the acetate concentration and the particles colored by the radius.

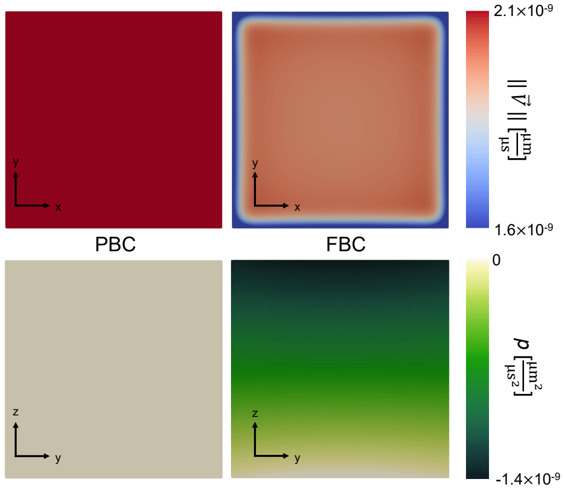


Figure S8: Comparing simulations with PBC and with FBC. Images of simulations with PBC (left) and with FBC (right). Simulations are colored by the magnitude of the fluid velocity ($\|\vec{V}\|$, top) or the kinematic pressure ($p$, bottom). These simulations used gravity, cells are hidden for clarity, and images are taken at an early timepoint (100 $\mu$s).

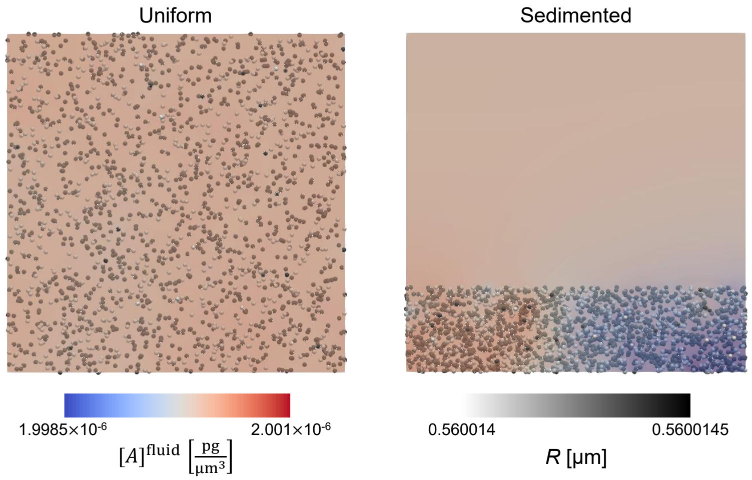


Figure S9: Comparison between simulations starting from uniform or sedimented starting positions. Images of simulations with uniform (left) and sedimented (right) simulations with the solution colored by the acetate concentration and the particles colored by the radius.

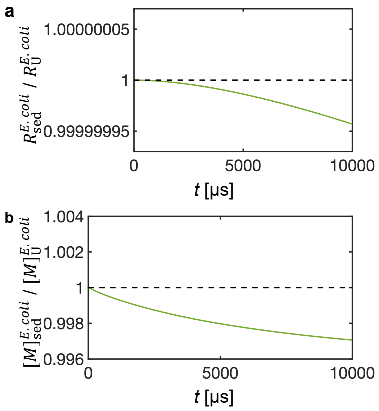


Figure S10: Comparison between simulations starting from uniform or sedimented starting positions with low metabolite concentration, gravity and FBC. The ratio between the average *E. coli* radius in sedimented ($R_{\rm sed}^{E.coli}$) and uniform ($R_{\rm U}^{E.coli}$) starting positions (a), and the ratio between the average methionine concentration in *E. coli* from sedimented ($[M]_{\rm sed}^{E.coli}$) and uniform ($[M]_{\rm U}^{E.coli}$) starting positions (b) as a function of time.

Table S1: Variables for the Monod cell growth. Square brackets are used to label mass units, i.e. pg[S] means pg of substrate.

| Variable | Symbol | Value at *E. coli* | Value at *S. enterica* |
|---|---|---|---|
| Maximum growth rate | $\mu_{\mathrm{max}}$ | $6.06 \times 10^{-8}\ \mu\mathrm{s}^{-1}$ | $3.03 \times 10^{-8}\ \mu\mathrm{s}^{-1}$ |
| Substrate concentration at which the growth rate is half of $\mu_{\mathrm{max}}$ | $K_{\mathrm{S}}$ | $2 \times 10^{-6}\ \frac{\mathrm{pg}}{\mu\mathrm{m}^3}$ | $1 \times 10^{-6}\ \frac{\mathrm{pg}}{\mu\mathrm{m}^3}$ |
| Growth yield coefficient | $Y_{\mathrm{MS}}$ | $0.5\frac{\mathrm{pg[M_i]}}{\mathrm{pg[S_i]}}$ | $0.5\frac{\mathrm{pg[M_i]}}{\mathrm{pg[S_i]}}$ |
| Yield of $P$ produced per $S$ consumed | $Y_{\mathrm{PS}}$ | $0.5\frac{\mathrm{pg[P_i]}}{\mathrm{pg[S_i]}}$ | $0.5\frac{\mathrm{pg[P_i]}}{\mathrm{pg[S_i]}}$ |

Table S2: Variables for the CFD portion of the simulation.

| Variable | Symbol | Value |
| --- | --- | --- |
| Time-step | $\delta t$ | $0.1\ \mu s$ |
| Simulation time | $t$ | $10^4\ \mu s$ |
| Interval frequency (for analysis) | $\Delta t$ | $10^2\ \mu s$ |
| Diffusion coefficient of methionine | $D_{met}$ | $1 \times 10^{-3}\ \frac{\mu m^2}{\mu s}$ |
| Diffusion coefficient of acetate | $D_{ace}$ | $1 \times 10^{-3}\ \frac{\mu m^2}{\mu s}$ |
| Volume of the simulation | $V_{sim}$ | $300\mu m \times 300\mu m \times 300\mu m$ |
| Volume of each uniform mesh bin | $V_{mesh}$ | $5\mu m \times 5\mu m \times 5\mu m$ |
| Initial kinematic pressure | $p_0$ | $0\ \frac{\mu m^2}{\mu s^2}$ |
| Initial velocity | $\vec{V}_0$ | $(0,\ 0,\ 0)\ \frac{\mu m}{\mu s}$ |
| Kinematic viscosity of the media | $\nu_l$ | $1.0\ \frac{\mu m^2}{\mu s}$ |

Table S3: Variables for the DEM portion of the simulation.

| Variable | Symbol | Value |
|---|---|---|
| Time-step | $\delta t$ | $0.1\ \mu$s |
| Simulation time | $t$ | $10^4\ \mu$s |
| Interval frequency (for analysis) | $\Delta t$ | $10^2\ \mu$s |
| Young's Modulus | $E$ | $2060\ \frac{pg}{\mu\text{m}\times\mu\text{s}^2}$ |
| Poisson Ratio | $\nu_\text{s}$ | 0.3 |
| Coefficient of restitution | $COR$ | 0.5 |
| Coefficient of friction | $COF$ | 0.0 |
| Cell density | $\rho$ | $1.08\ \frac{\text{pg}}{\mu\text{m}^3}$ |
| Initial radius | $r_0$ | $0.56\ \mu$m |

Table S4: Variables for the coupling the CFD and DEM portions of the simulation.

| Variable | Symbol | Value |
|---|---|---|
| Coupling interval | $\delta\delta t$ | 1 $\mu$s |
| Fluid thermal conductivity | $\lambda$ | 608.9 $\frac{\mathrm{pg}\times\mu\mathrm{m}}{\mu\mathrm{s}^3\mathrm{K}}$ |
| Prandtl number | Pr | 7.56 |
| Volumetric heat capacity of the liquid | $C_\mathrm{v}$ | 4180 $\frac{\mathrm{pg}}{\mu\mathrm{m}\times\mu\mathrm{s}^2 K}$ |
| Momentum exchange between the solid and liquid phases | $K_\mathrm{sl}$ | 0 $\frac{\mathrm{pg}}{\mu\mathrm{m}^3\mu\mathrm{s}}$ |
| Thermal conductivity of the particles | $k$ | 600 $\frac{\mathrm{pg}\times\mu\mathrm{m}}{\mu\mathrm{s}^3\mathrm{K}}$ |
| Thermal capacity of the particles | $C_\mathrm{s}$ | 4184 $\frac{\mu\mathrm{m}^2}{\mu\mathrm{s}^2 K}$ |

### Installation Guide

For any system, installation begins by downloading LIGGGHTS (https://github.com/CFDEMproject/LIGGGHTS-PUBLIC.git), OpenFOAM (https://github.com/OpenFOAM/OpenFOAM-5.x), and CFDEM®coupling (https://github.com/CFDEMproject/CFDEMcoupling-PUBLIC). Next, the user adds the CAMDLES2 source code to LIGGGHTS by copying `fix_cfd_coupling_convection_species.cpp`, `fix_cfd_coupling_convection_species.h`, `fix_scalar_transport_equation.cpp`, and `fix_scalar_transport_equation.h` to the user's `LIGGGHTS/LIGGGHTS-PUBLIC/src/`. Similarly, the user adds the CAMDLES2 source code to CFDEM®coupling by copying `cfdemSolverPisoSTM.C` to the user's `CFDEM/CFDEMcoupling-PUBLIC-5.x/applications/solvers/cfdemSolverPisoSTM/`. After these modifications, the user installs CFDEM®coupling according to standard procedures. More details are provided for specific systems below.

The installation for CAMDLES2 has been primarily tested for native Linux Ubuntu 18.04 via Bash. Compatible operating systems include:

- Ubuntu 18.04
- Red Hat Enterprise Linux (RHEL)
- Arch Linux
- Linux virtualization on macOS and Windows

Installation instructions for CFDEM®coupling have been adapted from https://www.engineerdo.com/wp-content/2021/02/Installation_CFDEM.pdf.

#### Ubuntu 18.04

1. Install core utilities:

   ```
   sudo apt update

   sudo upgrade

   sudo add-apt-repository main

   sudo add-apt-repository universe

   sudo apt-get install ffmpeg git-core
   ```

2. Clone the Git repositories for CFDEM® and LIGGGHTS®:

   ```
   cd $HOME

   mkdir CFDEM

   cd CFDEM

   git clone https://github.com/CFDEMproject/CFDEMcoupling-PUBLIC.git


   cd $HOME

   mkdir LIGGGHTS

   cd LIGGGHTS

   git clone https://github.com/CFDEMproject/LIGGGHTS-PUBLIC.git
   ```

3. Clone the Git repositories for OpenFOAM®:

   ```
   cd $HOME

   mkdir OpenFOAM

   git clone https://github.com/OpenFOAM/OpenFOAM-5.x.git

   git clone https://github.com/OpenFOAM/ThirdParty-5.x.git
   ```

4. Install essential packages:

sudo apt-get install build-essential flex bison cmake zlib1g-dev libboost-system-dev libboost-thread-dev libopenmpi-dev openmpi-bin gnuplot libreadline-dev libncurses-dev libxt-dev libscotch-dev libptscotch-dev python3-numpy python3-paraview

5. Edit and source ~/.bashrc:

   Add the following lines at the end of ~/.bashrc:

   ```
   export WM_NCOMPPROCS=12
   source $HOME/OpenFOAM/OpenFOAM-5.x/etc/bashrc
   ```

   Then save and exit. Then in the terminal:

   ```
   source ~/.bashrc
   ```

6. Compile OpenFOAM®:

   ```
   cd  /OpenFOAM/OpenFOAM-5.x/
   ./Allwmake
   ```

7. Add cell growth model and the CFD-DEM solver with gravity to CFDEM® and LIGGGHTS®. Download CAMDLES supplementary files, then:

   - Take the LIGGGHTS® files (`fix_cfd_coupling_convection_species.cpp`, `fix_cfd_coupling_convection_species.h`, `fix_scalar_transport_equation.cpp`, and `fix_scalar_transport_equation.h`) and add them to `LIGGGHTS-PUBLIC/src`.
   - Take `cfdemSolverPisoSTM.C` and add it to `CFDEMcoupling-PUBLIC/applications/solvers/cfdemSolverPisoSTM/`

8. Modify ~/.bashrc to source the CFDEM® directories:

```
cd $HOME/CFDEM
vim ~/.bashrc
```

and add the following lines to the end of the `~/.bashrc`:

```
# cfdem environment variables
export CFDEM_VERSION=PUBLIC
export CFDEM_PROJECT_DIR=$HOME/CFDEM/CFDEMcoupling-$CFDEM_VERSION-
$WM_PROJECT_VERSION
export CFDEM_PROJECT_USER_DIR=$HOME/CFDEM/$LOGNAME-
$CFDEM_VERSION-$WM_PROJECT_VERSION
export CFDEM_bashrc=$CFDEM_PROJECT_DIR/src/lagrangian/cfdemParticle/etc/bashrc
export CFDEM_LIGGGHTS_SRC_DIR=$HOME/LIGGGHTS/LIGGGHTS-PUBLIC/src
export CFDEM_LIGGGHTS_MAKEFILE_NAME=auto
.  $CFDEM_bashrc
```

9. Compile CFDEM®:

```
source ~/.bashrc
cfdemCompCFDEMall
```

10. Run the simulations (this may take >2 h) by downloading the code from *Example* and executing:

```
./Allrun.sh
```

#### Red Hat Enterprise Linux (RHEL)

This set of instructions is for running CAMDLES2 on the NASA Advanced Supercomputing (NAS) and other enterprise clusters running Red Hat Enterprise (RHEL) without sudo access. This instruction set also assumes use of the Portable Batch System (PBS) scheduler.

1. Navigate to your home directory (assumed here to be ∼) and download CFDEM®and LIGGGHTS®:

   ```
   cd ~
   mkdir CFDEM && cd CFDEM
   git clone https://github.com/CFDEMproject/CFDEMcoupling-PUBLIC.git

   cd ~
   mkdir LIGGGHTS && cd LIGGGHTS
   git clone https://github.com/CFDEMproject/LIGGGHTS-PUBLIC.git
   ```

2. Download OpenFOAM®:

   ```
   cd ~
   mkdir OpenFOAM && cd OpenFOAM
   git clone https://github.com/OpenFOAM/OpenFOAM-5.x.git
   git clone https://github.com/OpenFOAM/ThirdParty-5.x.git
   ```

3. Load in required modules for compilation using the module load command. These will vary based on specific HPC configuration. Line 89 in /OpenFOAM/OpenFOAM-5.x/etc/bashrc may need to be changed from "`export WM_MPLIB=SYSTEMOPENMPI`" to "`export WM_MPLIB=INTELMPI`". Modules may need to be reloaded if the session is exited.

4. The "scotch" package will likely not be available as a module in the HPC environment; install it with:

```
git clone https://gitlab.inria.fr/scotch/scotch && cd scotch
mkdir build && cd build
cmake ..  -DCMAKE_INSTALL_PREFIX=$HOME/.local
```

Note that the above flag is recommended because standard users usually do not have access to higher directories.

Then, line 5 in cmake_install.cmake should be changed from:

“`set (CMAKE_INSTALL_PREFIX ‘‘/usr/local’’)`” to “`set (CMAKE_INSTALL_PREFIX ‘‘$HOME/.local’’)`”

Then, install the “scotch” package with:

```
make && make install
```

5. Now, ensure that scotch, OpenFOAM, and MPI can be accessed on their respective paths. In ∼/.bashrc, add the following lines if they are not there already:

```
export PATH=$HOME/.local/bin:/<PATH_TO_MPI_SOURCE>/include:$PATH
export LD_LIBRARY_PATH=$HOME/.local/lib:$LD_LIBRARY_PATH
# OpenFOAM
export WM_NCOMPPROCS=<NUMBER_OF_AVAILABLE_PROCESSORS>
source $HOME/OpenFOAM/OpenFOAM-5.x/etc/bashrc
```

Save and exit, then

```
source ~/.bashrc
```

This should not return any message if everything was installed properly.

6. Add supplementary LIGGGHTS® and CFDEM®coupling files and test cases to the installation.

   - On a local machine, download the files and example included as text in this document and copy them into the cluster using the scp command or similar:

     ```
     scp -r ./<PATH_TO_SUPPLEMENTARY_FILES> <USERNAME>@<HOSTNAME>:~/
     ```

   - Take the LIGGGHTS® files (`fix_cfd_coupling_convection_species.cpp`, fix_cfd_coupling_convection_species.h,

fix_scalar_transport_equation.cpp,

and fix_scalar_transport_equation.h) and add them to LIGGGHTS-PUBLIC/src.

- Take cfdemSolverPisoSTM.C and add it to CFDEMcoupling-PUBLIC/applications/solvers/cfdemSolverPisoSTM/

7. Modify ~/.bashrc to source the CFDEM® directories by adding the following lines to the end of the file:

```
# cfdem environment variables
export CFDEM_VERSION=PUBLIC
export CFDEM_PROJECT_DIR=$HOME/CFDEM/CFDEMcoupling-$CFDEM_VERSION-
$WM_PROJECT_VERSION
export CFDEM_PROJECT_USER_DIR=$HOME/CFDEM/$LOGNAME-$CFDEM_VERSION-
$WM_PROJECT_VERSION
export CFDEM_bashrc=$CFDEM_PROJECT_DIR/src/lagrangian/cfdemParticle/etc/bashrc
export CFDEM_LIGGGHTS_SRC_DIR=$HOME/LIGGGHTS/LIGGGHTS-PUBLIC/src
export CFDEM_LIGGGHTS_MAKEFILE_NAME=auto
. $CFDEM_bashrc
```

Then source the bashrc file and compile CFDEM® using:

```
source ~/.bashrc
cfdemCompCFDEMall
```

8. Once CFDEM®coupling is successfully compiled, create a submission script called submission_script.sh. Here, is an example:

```
#PBS -S /bin/bash
#PBS -N CAMDLES
#PBS -l select=<SELECT>ncpus=<NCPUS>:model=<MODEL>
```

```
#PBS -q <QUEUE>
#PBS l walltime=<HH:MM:SS>
#PBS -W group_list=<GROUP_LIST>
module load <MODULE1>
module load <MODULE2>
./Allrun.sh > display
```

9. Note that the PBS submission script may need to be modified for each HPC. In particular, modifications to scripts may be need to efficiently parallelize the simulation among available processors:

   - In `parCFDDEMrun.sh`, change the number of processors (`nrProcs`) in line 22 to the number of available processors.
   - In `parDEMrun.sh`, change the number of processors (`nrProcs`) in line 23 to the number of available processors.
   - In `CFD/system/decomposeParDict`, change the number of subdomains (`numberofSubdomains`) in line 18 to the number of available processors.
   - In the same file, change n to a suitable number of partitions in the format ($x$ $y$ $z$), where $x$, $y$, and $z$ are positive integers and $x \times y \times z$ should equal the number of available processors.
   - in `DEM/in.liggghts_resume`, change the number of processor dimensions to $x$ $y$ $z$, constrained as above.

   To submit a job, first ensure that all environment variables are sourced

   ```
   source ~/.bashrc
   ```

   Then:

   ```
   qsub -V submission_script.sh
   ```

The -V flag submits the submission_script using the currently sourced environment variables. The job will enter the queue, and its status can be checked using:

```
qstat -nu <USERNAME>
```

To end a job early, type:

```
qdel <JOB_ID>
```

where <JOB_ID> is an eight-digit number shown in the first column of qstat.

#### Arch Linux - EndeavourOS

The installation of CAMDLES2 in Arch Linux is similar to that of Ubuntu, except the package manager "pacman" is used instead of apt-get to install the core utilities and essential packages. Note that some of the packages are already installed, some have simpler names, and some Arch Linux packages cover more than one dependency. To install the required packages:

```
sudo pacman -S ffmpeg git-core
sudo pacman -S cmake zlib boost openmpi gnuplot readline ncurses libxt python-numpy
paraview
```

The "flex" and "bison" packages will likely be installed; build-essential is only required in Ubuntu and Debian (for compiling). VTK should be a dependency of either Open MPI or Paraview.

However, the scotch package is not in the Arch Linux core repositories. It must be git cloned and cmake build modified with the position independent code flag:

```
cd $HOME
git clone https://gitlab.inria.fr/scotch/scotch
cd scotch
mkdir build && cd build
cmake ..  -DCMAKE_POSITION_INDEPENDENT_CODE=TRUE
```

```
make
```

make install

#### Linux virtualization on macOS and Windows

Many dependencies of CAMDLES2 are not currently compatible with macOS and Windows operating systems and virtualization software should be used. Please allocate ∼8-10 GB RAM and >64 GB of storage memory to run properly. For users with Apple Silicon, ensure that the virtualization software allows the CAMDLES2 installation to take advantage of hardware acceleration.

### Supplemental Code

#### Source Code

Usage:

- To enable Monod-growth based on metabolite consumption, metabolites must be chemical species in the solution. Thus, input files specifying initial concentrations should exist in `CFD/0`.

- Further, each metabolite should be referenced in `CFD/constant/couplingProperties`, which should include `scalarGeneralExchange` as a force model. Each metabolite should be referenced in `scalarGeneralExchangeProps` in `partSpeciesNames`

- Next, to execute the growth reaction, the reaction command must be executed in the DEM portion of the simulation. Specifically, in `in.liggghts_resume`, a `fix couple/cfd/speciesConvection` call should be executed. This call executes convection between the biological cells and the fluid, and calls fix scalar transport equation, which executes the Monod-based cell growth.

- The syntax of `fix couple/cfd/speciesConvection` is as follows ({brackets} indicates that these are the values of the specified keyword):
`fix FixName all couple/cfd/speciesConvection speciesName {speciesName} species0 {species0} yield_product_from_substrate {yield_product_from_substrate} atomtype {atomtype} SubstrateName {SubstrateName} u_max {u_max} K_S {K_S} yield_growth_coefficient {yield_growth_coefficient}`
where {`speciesName`} is the species that is produced by the reaction, {`species0`} is the initial concentration of {speciesName}, {`yield_product_from_substrate`} is the mass of {speciesName} created by the reaction per mass of {SubstrateName}, {`atomtype`} is the atomtype that produces {speciesName}, {`SubstrateName`} is the species consumed by the reaction, {`u_max`} is the maximum growth rate for the microbe, {`K_S`} is

the half-saturation constant (substrate concentration at which the growth rate is half of its maximum value), and {`yield_growth_coefficient`} is the amount of biomass produced per unit of substrate consumed

Summary of installation:

1. Download LIGGGHTS (https://github.com/CFDEMproject/LIGGGHTS-PUBLIC.git)
2. Download OpenFOAM (https://github.com/OpenFOAM/OpenFOAM-5.x)
3. Download CFDEM (https://github.com/CFDEMproject/CFDEMcoupling-PUBLIC)
4. Replace `fix_cfd_coupling_convection_species.cpp`, `fix_cfd_coupling_convection_species.h`, `fix_scalar_transport_equation.cpp`, and `fix_scalar_transport_equation.h` in your version of `LIGGGHTS/LIGGGHTS-PUBLIC/src/` with those from this file.
5. Replace `cfdemSolverPisoSTM.C` in your version of `CFDEM/CFDEMcoupling-PUBLIC-5.x/applications/solvers/cfdemSolverPisoSTM/` with that from this file.
6. Download and install VTK (https://github.com/Kitware/VTK.git).
7. Download and install scotch (https://gitlab.inria.fr/scotch/scotch).
8. Compile OpenFOAM.
9. Compile CFDEM, which will compile LIGGGHTS by default.

For additional help with steps 6-9, see:
`https://www.cfdem.com/media/CFDEM/docu/CFDEMcoupling_Manual.html#installation` and `https://www.youtube.com/watch?v=V6VaVE6VB0g&t=700s`.

**fix_cfd_coupling_convection_species.cpp**

```
/* ----------------------------------------------------------------------
This is the LIGGGHTS
DEM simulation engine, released by
DCS Computing Gmbh, Linz, Austria
http://www.dcs-computing.com,

LIGGGHTS® is part of CFDEM®project:
http://www.liggghts.com | http://www.cfdem.com

Core developer and main author:
Christoph Kloss,
Modified for biological cell growth by:
Andrew P. Latham, Emmanuel N. Skountzos, Tyler Quarton,
and Rocky An

LIGGGHTS® is open-source, distributed under the terms of the GNU Public
License, version 2 or later. It is distributed in the hope that it will
be useful, but WITHOUT ANY WARRANTY; without even the implied warranty
of MERCHANTABILITY or FITNESS FOR A PARTICULAR PURPOSE. You should have
received a copy of the GNU General Public License along with LIGGGHTS®.
If not, see http://www.gnu.org/licenses . See also top-level README
and LICENSE files.

LIGGGHTS® and CFDEM® are registered trade marks of DCS Computing GmbH,
the producer of the LIGGGHTS® software and the CFDEM®coupling software
See http://www.cfdem.com/terms-trademark-policy for details.
```

```
------------------------------------------------------------------------
   Contributing author and copyright for this file:
   (if not contributing author is listed, this file has been contributed
   by the core developer)

   Copyright 2015-     DCS Computing GmbH, Linz
------------------------------------------------------------------------- */

#include <string.h>
#include <stdlib.h>
#include "atom.h"
#include "update.h"
#include "respa.h"
#include "error.h"
#include "neighbor.h"
#include "memory.h"
#include "modify.h"
#include "group.h"
#include "comm.h"
#include <cmath>
#include "vector_liggghts.h"
#include "fix_cfd_coupling_convection_species.h"
#include "fix_property_atom.h"
#include "fix_property_global.h"
#include "properties.h"
```

```
using namespace LAMMPS_NS;
using namespace FixConst;

/* ---------------------------------------------------------------------- */

// fix that executes species transfer between the particles and the fluid. Here, this
// function has been modified to execute particle (biological cell) growth in addition
// to species transfer.
// Growth depends on one species (called SubstrateName) and produces a second species
// (called speciesName)
FixCfdCouplingConvectionSpecies::FixCfdCouplingConvectionSpecies(LAMMPS *lmp,
    int narg, char **arg) :  Fix(lmp, narg, arg)
{
    fix_coupling = NULL;
    fix_speciesConcentration = fix_convectiveFlux = fix_totalFlux = NULL;
    fix_speciesFluid = fix_speciesTransCoeff = NULL;

    // 16 Arguments. Each is specified below
    int iarg = 3;

    if(narg < iarg + 16) error->all(FLERR,
    "Fix couple/cfd/speciesConvection: Wrong number of arguments. Expected format:
    {fix_name} all couple/cfd/speciesConvection speciesName {speciesName} species0
    {species0} yield_product_from_substrate {yield_product_from_substrate} atomtype
    {atomtype} SubstrateName {SubstrateName} u_max {u_max} K_S {K_S}
    yield_growth_coefficient {yield_growth_coefficient}");
    // Argument 1 - speciesName. This should correspond to the
```

```
// species that is produced by the reaction.
if(strcmp(arg[iarg++],"speciesName") != 0) error->all(FLERR,"Fix
        couple/cfd/speciesConvection: Expecting keyword 'speciesName'");
    strcpy(speciesName_,arg[iarg++]);
    // These lines initialize the name of other variables
    sprintf(sourceName_,        "%sSource",speciesName_);
    sprintf(convectiveFluxName_,"%sFlux",speciesName_);
    sprintf(capacityName_,      "%sCapacity",speciesName_);
    sprintf(steName_,           "%sSTE",speciesName_);
    sprintf(totalFluxName_,     "%sTotalFlux",speciesName_);
    sprintf(speciesFluidName_,      "%sFluid",speciesName_);
    sprintf(speciesTransCoeffName_, "%sTransCoeff",speciesName_);

// Argument 2/3 - species0 keyword, followed by value.
if(strcmp(arg[iarg++],"species0") != 0) error->all(FLERR,
    "Fix couple/cfd/speciesConvection: Expecting keyword 'species0'");
    species0 = atof(arg[iarg++]);
// Check initial species0
if(species0 < 0.) error->all(FLERR,
    "Fix couple/cfd/speciesConvection: species0 must be >= 0");

// Argument 4/5 - yield_product_from_substrate keyword, followed by value.
// This is the mass of speciesName created by the reaction per mass
// of SubstrateName
if(strcmp(arg[iarg++],"yield_product_from_substrate") != 0) error->all(FLERR,
        "Fix couple/cfd/speciesConvection: Expecting keyword
        'yield_product_from_substrate'");
```

```
    yield_product_from_substrate = atof(arg[iarg++]);

// Argument 6/7 - atomtype keyword, followed by value.
// This is the atomtype that produces speciesName
if(strcmp(arg[iarg++],"atomtype") != 0) error->all(FLERR,
        "Fix couple/cfd/speciesConvection: Expecting keyword 'atomtype'");
    atomtype = std::stoi(arg[iarg++]);

// Argument 8/9 - SubstrateName keyword, followed by value.
// This is the species consumed by the reaction
if(strcmp(arg[iarg++],"SubstrateName") != 0) error->all(FLERR,
        "Fix couple/cfd/speciesConvection: Expecting keyword 'SubstrateName'");
    strcpy(SubstrateName,arg[iarg++]);

// Argument 10/11 - u_max keyword, followed by value.
// u_max is the maximum growth rate for the microorganism,
// specified by the atomtype
if(strcmp(arg[iarg++],"u_max") != 0) error->all(FLERR,
        "Fix couple/cfd/speciesConvection: Expecting keyword 'u_max'");
    u_max = atof(arg[iarg++]);

// Argument 12/13 - K_S keyword, followed by value.
// K_S is the half-saturation constant
// (substrate concentration at which the growth rate is half of its maximum value)
if(strcmp(arg[iarg++],"K_S") != 0) error->all(FLERR,
        "Fix couple/cfd/speciesConvection: Expecting keyword 'K_S'");
    K_S = atof(arg[iarg++]);
```

```
    // Argument 14/15 - yield_growth_coefficient keyword, followed by value.
    // This is the growth yield coefficient, which represents
    // the amount of biomass produced per unit of substrate consumed
    if(strcmp(arg[iarg++],"yield_growth_coefficient") != 0) error->all(FLERR,
            "Fix couple/cfd/speciesConvection: Expecting keyword
            'yield_growth_coefficient'");
        yield_growth_coefficient = atof(arg[iarg++]);

}

/* ---------------------------------------------------------------------- */
FixCfdCouplingConvectionSpecies::~FixCfdCouplingConvectionSpecies()
{
}

/* ---------------------------------------------------------------------- */

void FixCfdCouplingConvectionSpecies::pre_delete(bool unfixflag)
{
    if(fix_convectiveFlux) modify->delete_fix(convectiveFluxName_);
}

/* ---------------------------------------------------------------------- */

int FixCfdCouplingConvectionSpecies::setmask()
{
```

```
  int mask = 0;
  mask |= POST_FORCE;
  return mask;
}

/* ----------------------------------------------------------------------- */

void FixCfdCouplingConvectionSpecies::post_create()
{

  //  register species concentration
  if(!fix_speciesConcentration)
  {
        const char* fixarg[9];
        fixarg[0]=speciesName_;
        fixarg[1]="all";
        fixarg[2]="property/atom";
        fixarg[3]=speciesName_;
        fixarg[4]="scalar";
        fixarg[5]="no";
        fixarg[6]="yes";
        fixarg[7]="no";
        fixarg[8]="0.";
        fix_speciesConcentration = modify->
            add_fix_property_atom(9,const_cast<char**>(fixarg),style);
  }
```

```
//  register convective flux
if(!fix_convectiveFlux)
{
      const char* fixarg[9];
      fixarg[0]=convectiveFluxName_;
      fixarg[1]="all";
      fixarg[2]="property/atom";
      fixarg[3]=convectiveFluxName_;
      fixarg[4]="scalar";
      fixarg[5]="no";
      fixarg[6]="yes";
      fixarg[7]="no";
      fixarg[8]="0.";
      fix_convectiveFlux = modify->
          add_fix_property_atom(9,const_cast<char**>(fixarg),style);
}

//  add species transfer model if not yet active
if(!fix_speciesFluid)
{
      const char* fixarg[9];
      fixarg[0]=speciesFluidName_;
      fixarg[1]="all";
      fixarg[2]="property/atom";
      fixarg[3]=speciesFluidName_;
      fixarg[4]="scalar";
      fixarg[5]="no";
```

```
        fixarg[6]="yes";
        fixarg[7]="no";
        fixarg[8]="0.";
        fix_speciesFluid = modify->
            add_fix_property_atom(9,const_cast<char**>(fixarg),style);
}

//  register transfer coefficient
if(!fix_speciesTransCoeff)
{
        const char* fixarg[9];
        fixarg[0]=speciesTransCoeffName_;
        fixarg[1]="all";
        fixarg[2]="property/atom";
        fixarg[3]=speciesTransCoeffName_;
        fixarg[4]="scalar";
        fixarg[5]="no";
        fixarg[6]="yes";
        fixarg[7]="no";
        fixarg[8]="0.";
        fix_speciesTransCoeff = modify->
            add_fix_property_atom(9,const_cast<char**>(fixarg),style);
}
// FixScalarTransportEquation - this fix executes cell growth
FixScalarTransportEquation *fix_ste =
modify->find_fix_scalar_transport_equation("speciesTransfer");
if(!fix_ste)
```

```
{
      int max_type = atom->get_properties()->max_type();
      Fix* capacityQty = modify->find_fix_property(
          capacityName_,"property/global","peratomtype",max_type,0,style,false);

      if(capacityQty==NULL)
      {
          sprintf(capacityName_, "none");
          if(comm->me==0)
                printf("WARNING: FixCfdCouplingConvectionSpecies cannot locate
                capacity quantity. Thus, will assume you are not using capacity
                in scalar transport equation is '%s'.", capacityName_);

      }
      // Pass arguments to FixScalarTransportEquation. Include new arguments
      const char *newarg[27];
      newarg[0] = steName_;
      newarg[1] = group->names[igroup];
      newarg[2] = "transportequation/scalar";
      newarg[3] = "equation_id";
      newarg[4] = steName_;
      newarg[5] = "quantity";
      newarg[6] = speciesName_;
      newarg[7] = "default_value";
      // convert to string before input
      char arg8[30];
      sprintf(arg8,"%.20f",species0);
```

```
newarg[8] = arg8;
newarg[9]  = "flux_quantity";
newarg[10] = totalFluxName_;
newarg[11] = "source_quantity";
newarg[12] = sourceName_;
newarg[13] = "capacity_quantity";
newarg[14] = capacityName_;
newarg[15] = "yield_product_from_substrate";
// convert to string before input
char arg16[30];
sprintf(arg16,"%.20f",yield_product_from_substrate);
newarg[16] = arg16;
newarg[17] = "atomtype";
// convert to string before input
char arg18[30];
sprintf(arg18,"%i",atomtype);
newarg[18] = arg18;
newarg[19] = "SubstrateName";
newarg[20] = SubstrateName;
newarg[21] = "u_max";
// convert to string before input
char arg22[30];
sprintf(arg22,"%.20f",u_max);
newarg[22] = arg22;
newarg[23] = "K_S";
// convert to string before input
char arg24[30];
```

```
        sprintf(arg24,"%.20f",K_S);
        newarg[24] = arg24;
        newarg[25] = "yield_growth_coefficient";
        // convert to string before input
        char arg26[30];
        sprintf(arg26,"%.20f",yield_growth_coefficient);
        newarg[26] = arg26;
        modify->add_fix(27,const_cast<char**>(newarg));
  }
}

/* ---------------------------------------------------------------------- */

void FixCfdCouplingConvectionSpecies::init()
{
    // make sure there is only one fix of this style
    //if(modify->n_fixes_style(style) != 1)
    //  error->fix_error(FLERR,this,"More than one fix of this style is not allowed");

    // find coupling fix
    fix_coupling = static_cast<FixCfdCoupling*>
    (modify->find_fix_style_strict("couple/cfd",0));
    if(!fix_coupling)
      error->fix_error(FLERR,this,"needs a fix of type couple/cfd");

    //values to send to OF
    fix_coupling->add_push_property(speciesName_,"scalar-atom");
```

```
//values to come from OF
fix_coupling->add_pull_property(convectiveFluxName_,"scalar-atom");
fix_coupling->add_pull_property(sourceName_,"scalar-atom");

fix_coupling->add_pull_property(speciesFluidName_,"scalar-atom");
fix_coupling->add_pull_property(speciesTransCoeffName_,"scalar-atom");

fix_speciesConcentration  = static_cast<FixPropertyAtom*>
    (modify->find_fix_property(speciesName_,"property/atom",
    "scalar",0,0,style));
fix_convectiveFlux        = static_cast<FixPropertyAtom*>
    (modify->find_fix_property(convectiveFluxName_,
    "property/atom","scalar",0,0,style));
fix_totalFlux             = static_cast<FixPropertyAtom*>
    (modify->find_fix_property(totalFluxName_,"property/atom",
    "scalar",0,0,style));
fix_speciesFluid          = static_cast<FixPropertyAtom*>
    (modify->find_fix_property(speciesFluidName_,"property/atom",
    "scalar",0,0,style));
fix_speciesTransCoeff     = static_cast<FixPropertyAtom*>
    (modify->find_fix_property(speciesTransCoeffName_,
    "property/atom","scalar",0,0,style));

if(!fix_speciesConcentration || !fix_convectiveFlux)
  error->fix_error(FLERR,this,"could not find concentration and flux fix");
```

```
}

/* ---------------------------------------------------------------------- */

void FixCfdCouplingConvectionSpecies::post_force(int)
{

  int *mask = atom->mask;
  int nlocal = atom->nlocal;

  // communicate convective flux to ghosts, there might be new data
  if(0 == neighbor->ago)
        fix_convectiveFlux->do_forward_comm();

  double *totalFlux      = fix_totalFlux->vector_atom;
  double *convectiveFlux = fix_convectiveFlux->vector_atom;

  for (int i = 0; i < nlocal; i++)
    if (mask[i] & groupbit)
      totalFlux[i] += convectiveFlux[i];
}
```

**fix_cfd_coupling_convection_species.h**

```
/* ----------------------------------------------------------------------
    This is the LIGGGHTS
    DEM simulation engine, released by
    DCS Computing Gmbh, Linz, Austria
    http://www.dcs-computing.com,

    LIGGGHTS® is part of CFDEM®project:
    http://www.liggghts.com | http://www.cfdem.com

    Core developer and main author:
    Christoph Kloss,
    Modified for biological cell growth by:
    Andrew P. Latham, Emmanuel N. Skountzos, Tyler Quarton,
    and Rocky An

    LIGGGHTS® is open-source, distributed under the terms of the GNU Public
    License, version 2 or later. It is distributed in the hope that it will
    be useful, but WITHOUT ANY WARRANTY; without even the implied warranty
    of MERCHANTABILITY or FITNESS FOR A PARTICULAR PURPOSE. You should have
    received a copy of the GNU General Public License along with LIGGGHTS®.
    If not, see http://www.gnu.org/licenses . See also top-level README
    and LICENSE files.

    LIGGGHTS® and CFDEM® are registered trade marks of DCS Computing GmbH,
    the producer of the LIGGGHTS® software and the CFDEM®coupling software
    See http://www.cfdem.com/terms-trademark-policy for details.
```

```
------------------------------------------------------------------------
    Contributing author and copyright for this file:
    (if not contributing author is listed, this file has been contributed
    by the core developer)

    Copyright 2015-     DCS Computing GmbH, Linz
------------------------------------------------------------------------- */

#ifdef FIX_CLASS

FixStyle(couple/cfd/speciesConvection,FixCfdCouplingConvectionSpecies)

#else

#ifndef LMP_FIX_CFD_COUPLING_CONVECTION_SPECIES_H
#define LMP_FIX_CFD_COUPLING_CONVECTION_SPECIES_H

#include "fix_cfd_coupling.h"

namespace LAMMPS_NS {

class FixCfdCouplingConvectionSpecies : public Fix {

 public:
  FixCfdCouplingConvectionSpecies(class LAMMPS *, int, char **);
  ~FixCfdCouplingConvectionSpecies();
```

```
 void post_create();
 void pre_delete(bool unfixflag);

 virtual int  setmask();
 virtual void init();
 virtual void post_force(int);

protected:
 class FixCfdCoupling*  fix_coupling;
 class FixPropertyAtom* fix_speciesConcentration;
 class FixPropertyAtom* fix_speciesFluid;
 class FixPropertyAtom* fix_speciesTransCoeff;
 class FixPropertyAtom* fix_convectiveFlux;
 class FixPropertyAtom* fix_totalFlux;

 double species0;
 char   speciesName_[128];
 char   sourceName_[128];
 char   convectiveFluxName_[128];
 char   capacityName_[128];
 char   steName_[128];
 char   totalFluxName_[128];
 char   speciesFluidName_[128];
 char   speciesTransCoeffName_[128];
 // Added variables for biological cell growth
 double yield_product_from_substrate;
 double yield_growth_coefficient;
```

```
  int atomtype;
  char   SubstrateName[128];
  double u_max;
  double K_S;
};

}

#endif
#endif
```

**fix_scalar_transport_equation.cpp**

```
/* ----------------------------------------------------------------------
    This is the LIGGGHTS
    DEM simulation engine, released by
    DCS Computing Gmbh, Linz, Austria
    http://www.dcs-computing.com,

    LIGGGHTS® is part of CFDEM®project:
    http://www.liggghts.com | http://www.cfdem.com

    Core developer and main author:
    Christoph Kloss,
    Modified for biological cell growth by:
    Andrew P. Latham, Emmanuel N. Skountzos, Tyler Quarton,
    and Rocky An

    LIGGGHTS® is open-source, distributed under the terms of the GNU Public
    License, version 2 or later. It is distributed in the hope that it will
    be useful, but WITHOUT ANY WARRANTY; without even the implied warranty
    of MERCHANTABILITY or FITNESS FOR A PARTICULAR PURPOSE. You should have
    received a copy of the GNU General Public License along with LIGGGHTS®.
    If not, see http://www.gnu.org/licenses . See also top-level README
    and LICENSE files.

    LIGGGHTS® and CFDEM® are registered trade marks of DCS Computing GmbH,
    the producer of the LIGGGHTS® software and the CFDEM®coupling software
    See http://www.cfdem.com/terms-trademark-policy for details.
```

```
------------------------------------------------------------------------

    Contributing author and copyright for this file:
    (if not contributing author is listed, this file has been contributed
    by the core developer)

    Copyright 2012-     DCS Computing GmbH, Linz
    Copyright 2009-2012 JKU Linz

    Implementation of implicit update algorithm
    Copyright 2016      TU Graz, Stefan Radl
    Copyright 2016      DCS Computing GmbH, Linz
------------------------------------------------------------------------- */

#include <cmath>
#include <stdlib.h>
#include <string.h>
#include "fix_scalar_transport_equation.h"
#include "atom.h"
#include "domain.h"
#include "group.h"
#include "force.h"
#include "update.h"
#include "error.h"
#include "modify.h"
#include "neighbor.h"
#include "neigh_list.h"
```

```
#include "neigh_request.h"
#include "pair.h"
#include "math_extra.h"
#include "fix_property_global.h"
#include "fix_property_atom.h"
#include "respa.h"
#include "properties.h"
#include "pair_gran.h"
#include "mpi_liggghts.h"

#include "math_const.h"

using namespace MathConst;
using namespace LAMMPS_NS;
using namespace FixConst;

#define SMALL 1e-8

/* ---------------------------------------------------------------------- */

// Executes fixscalar transport equation. If the number of arguments is 27,
// the code assumes a biological cell growth reaction will occur.
// If the number of arguments is not 27, then the code executes
// normal scalar transport.
FixScalarTransportEquation::FixScalarTransportEquation(
    LAMMPS *lmp, int narg, char **arg) : Fix(lmp, narg, arg)
{
```

```
fix_quantity = fix_flux = fix_source = NULL; fix_capacity = NULL;
fix_fluidQty_ = fix_transCoeffQty_ = NULL;
fluidQty_0_       = 0;
transCoeffQty_0_  = 0;

capacity = NULL;

int_flag = true;

nevery_  = 1;
//ensure flux is reset at the very first time step
performedIntegrationLastStep_ = true;

peratom_flag = 1;
size_peratom_cols = 0;
peratom_freq = 1;

scalar_flag = 1;
global_freq = 1;

capacity_name = NULL;
capacity_flag = 0;

reaction_flag = false;
// If narg==27, then reaction_flag, which means FixScalarTransportEquation was
// called by FixCfdCouplingConvectionSpecies instead of by another function.
// This implies extra arguments are expected, which dictates a bioreaction
```

```
if(narg==27){
  reaction_flag = true;
  /*for (int i = 0; i < narg; i++){
    printf("arg: '%i'.\n", i);
    printf("arg: '%s'.\n", arg[i]);
  } */
}

crankNicholsonFactor_ = 0.5; //0 ... fully explicit; 1 ... fully implicit
implicitMode_ = false;
advanceQty=&FixScalarTransportEquation::advanceQtyExplicit;

if(strcmp(arg[2],"transportequation/scalar"))
  return;

// 25 args. Start at arg 3 to read in
int iarg = 3;
if (narg < 15)
  error->fix_error(FLERR,this,"not enough arguments");

// 3/4 equation_id, keyword and value
if(strcmp(arg[iarg++],"equation_id"))
  error->fix_error(FLERR,this,"expecting keyword 'equation_id'");
equation_id = new char[strlen(arg[iarg])+1];
strcpy(equation_id,arg[iarg++]);

// 5/6 quantity (speciesName_), keyword and value
```

```
if(strcmp(arg[iarg++],"quantity"))
  error->fix_error(FLERR,this,"expecting keyword 'quantity'");
quantity_name = new char[strlen(arg[iarg])+1];
strcpy(quantity_name,arg[iarg++]);

// 7/8 default_value (species0), keyword and value
if(strcmp(arg[iarg++],"default_value"))
  error->fix_error(FLERR,this,"expecting keyword 'default_value'");
quantity_0 = atof(arg[iarg++]);

// 9/10 flux_quantity (totalFluxName_), keyword and value
if(strcmp(arg[iarg++],"flux_quantity"))
  error->fix_error(FLERR,this,"expecting keyword 'flux_quantity'");
flux_name = new char[strlen(arg[iarg])+1];
strcpy(flux_name,arg[iarg++]);

// 11/12 source_quantity (sourceName_), keyword and value
if(strcmp(arg[iarg++],"source_quantity"))
  error->fix_error(FLERR,this,"expecting keyword 'source_quantity'");
source_name = new char[strlen(arg[iarg])+1];
strcpy(source_name,arg[iarg++]);

// 13/14 capacity_quantity (capacityName_), keyword and value
if(strcmp(arg[iarg++],"capacity_quantity"))
  error->fix_error(FLERR,this,"expecting keyword 'capacity_quantity'");
if(strcmp(arg[iarg],"none"))
{
```

```
    capacity_flag = 1;
    capacity_name = new char[strlen(arg[iarg])+1];
    strcpy(capacity_name,arg[iarg++]);
// make sure iarg iterates regardless
} else {
  iarg++;
}



// if reaction_flag, expect arguments related to the bioreaction
if (reaction_flag){
  // 15/16 yield_product_from_substrate (yield_product_from_substrate),
  // keyword and value
  if(strcmp(arg[iarg++],"yield_product_from_substrate"))
    error->fix_error(FLERR,this,"expecting keyword 'yield_product_from_substrate'");
  yield_product_from_substrate = atof(arg[iarg++]);

  // 17/18 atomtype (atomtype), keyword and value
  if(strcmp(arg[iarg++],"atomtype"))
    error->fix_error(FLERR,this,"expecting keyword 'atomtype'");
  atomtype = std::stoi(arg[iarg++]);

  // 19/20 SubstrateName (SubstrateName), keyword and value
  if(strcmp(arg[iarg++],"SubstrateName"))
    error->fix_error(FLERR,this,"expecting keyword 'SubstrateName'");
  SubstrateName = new char[strlen(arg[iarg])+1];
  strcpy(SubstrateName,arg[iarg++]);
```

```
    // 21/22 u_max (u_max), keyword and value
    if(strcmp(arg[iarg++],"u_max"))
      error->fix_error(FLERR,this,"expecting keyword 'u_max'");
    u_max = atof(arg[iarg++]);

    // 23/24 K_S (K_S), keyword and value
    if(strcmp(arg[iarg++],"K_S"))
      error->fix_error(FLERR,this,"expecting keyword 'K_S'");
    K_S = atof(arg[iarg++]);

    // 25/26 yield_growth_coefficient (yield_growth_coefficient), keyword and value
    if(strcmp(arg[iarg++],"yield_growth_coefficient"))
      error->fix_error(FLERR,this,"expecting keyword 'yield_growth_coefficient'");
    yield_growth_coefficient = atof(arg[iarg++]);
  }
}

/* ---------------------------------------------------------------------- */

FixScalarTransportEquation::~FixScalarTransportEquation()
{
    delete []quantity_name;
    delete []flux_name;
    delete []source_name;
    delete []capacity_name;
    delete []equation_id;
```

```
    if(implicitMode_)
    {
        delete []fluid_name_;
        delete []transCoeff_name_;
    }

    if(capacity) delete []capacity;

}

/* ---------------------------------------------------------------------- */
void FixScalarTransportEquation::register_implicit_fixes(
    char* fluidName, double fluid0, char* transCoeffName, double transCoeff0)
{
  const char *fixarg[9];

  implicitMode_ = true;
  advanceQty=&FixScalarTransportEquation::advanceQtyImplicit;

  fluid_name_ = new char[strlen(fluidName)+1];
  strcpy(fluid_name_,fluidName);
  fluidQty_0_ = fluid0;

  transCoeff_name_ = new char[strlen(transCoeffName)+1];
  strcpy(transCoeff_name_,transCoeffName);
  transCoeffQty_0_ = transCoeff0;
```

```
fix_fluidQty_=static_cast<FixPropertyAtom*>(
  modify->find_fix_property(fluid_name_,"property/atom","scalar",0,0,style));
if (fix_fluidQty_==NULL) {
      //register fluid quantity as property/atom
      fixarg[0]=fluid_name_;
      fixarg[1]="all";
      fixarg[2]="property/atom";
      fixarg[3]=fluid_name_;
      fixarg[4]="scalar";
      fixarg[5]="yes";    //restart
      fixarg[6]="no";     //commGhost
      fixarg[7]="yes";    //commGhostRev
      char arg8[30];
      sprintf(arg8,"%e", fluidQty_0_);
      fixarg[8]=arg8;
      modify->add_fix(9,const_cast<char**>(fixarg));
      fix_fluidQty_=static_cast<FixPropertyAtom*>(
          modify->find_fix_property(fluid_name_,"property/atom",
          "scalar",0,0,style));
}

fix_transCoeffQty_=static_cast<FixPropertyAtom*>(
  modify->find_fix_property(transCoeff_name_,"property/atom","scalar",0,0,style));
if (fix_transCoeffQty_==NULL){
      //register transfer coefficient as property/atom
      fixarg[0]=transCoeff_name_;
```

```
        fixarg[1]="all";
        fixarg[2]="property/atom";
        fixarg[3]=transCoeff_name_;
        fixarg[4]="scalar";
        fixarg[5]="yes";    //restart
        fixarg[6]="no";     //commGhost
        fixarg[7]="yes";    //commGhostRev
        char arg8[30];
        sprintf(arg8,"%e", transCoeffQty_0_);
        fixarg[8]=arg8;
        modify->add_fix(9,const_cast<char**>(fixarg));
        fix_transCoeffQty_=static_cast<FixPropertyAtom*>(
            modify->find_fix_property(transCoeff_name_,"property/atom",
            "scalar",0,0,style));
  }

  updatePtrsImpl();

  return;
}

/* ---------------------------------------------------------------------- */

void FixScalarTransportEquation::pre_delete(bool unfixflag)
{
    //unregister property/atom fixes
    if(unfixflag)
```

```
    {
        if (fix_quantity) modify->delete_fix(quantity_name);
        if (fix_flux) modify->delete_fix(flux_name);
        if (fix_source) modify->delete_fix(source_name);
        if(implicitMode_)
        {
            if (fix_fluidQty_) modify->delete_fix(fluid_name_);
            if (fix_transCoeffQty_) modify->delete_fix(transCoeff_name_);
        }
    }
}

/* ---------------------------------------------------------------------- */

int FixScalarTransportEquation::setmask()
{
  int mask = 0;
  mask |= INITIAL_INTEGRATE_RESPA;
  mask |= INITIAL_INTEGRATE;
  mask |= PRE_FORCE;
  mask |= FINAL_INTEGRATE;
  return mask;
}

/* ---------------------------------------------------------------------- */

void FixScalarTransportEquation::updatePtrs()
```

```
{
  quantity = fix_quantity->vector_atom;
  flux = fix_flux->vector_atom;
  source = fix_source->vector_atom;

  if(implicitMode_)
    updatePtrsImpl();

  vector_atom = quantity;
}

/* ---------------------------------------------------------------------- */
void FixScalarTransportEquation::updatePtrsImpl()
{
  fluidQty_         = fix_fluidQty_->vector_atom;
  transCoeffQty_    = fix_transCoeffQty_->vector_atom;
}

/* ---------------------------------------------------------------------- */

void FixScalarTransportEquation::post_create()
{
  const char *fixarg[9];

  if (fix_quantity==NULL) {
    //register Temp as property/atom
    // For biological contexts, the quantity is the
```

```
  // biological concentration of a metabolite
  fixarg[0]=quantity_name;
  fixarg[1]="all";
  fixarg[2]="property/atom";
  fixarg[3]=quantity_name;
  fixarg[4]="scalar";
  fixarg[5]="yes";
  fixarg[6]="yes";
  fixarg[7]="no";
  char arg8[30];
  sprintf(arg8,"%e",quantity_0);
  fixarg[8]=arg8;
  modify->add_fix(9,const_cast<char**>(fixarg));
  fix_quantity=static_cast<FixPropertyAtom*>(
      modify->find_fix_property(quantity_name,"property/atom",
      "scalar",0,0,style));
}

if (fix_flux==NULL){
  //register heatFlux as property/atom
  // For biological contexts, flux is the
  // inward flux of metabolite from the solution.
  fixarg[0]=flux_name;
  fixarg[1]="all";
  fixarg[2]="property/atom";
  fixarg[3]=flux_name;
  fixarg[4]="scalar";
```

```
    fixarg[5]="yes";
    fixarg[6]="no";
    fixarg[7]="yes";
    fixarg[8]="0.";
    modify->add_fix(9,const_cast<char**>(fixarg));
    fix_flux=static_cast<FixPropertyAtom*>(
        modify->find_fix_property(flux_name,"property/atom","scalar",0,0,style));
  }

  if (fix_source==NULL){
    //register heatSource as property/atom
    // For biological contexts, source is the
    // produced metabolite excreted to the solution.
    fixarg[0]=source_name;
    fixarg[1]="all";
    fixarg[2]="property/atom";
    fixarg[3]=source_name;
    fixarg[4]="scalar";
    fixarg[5]="yes";
    fixarg[6]="yes";
    fixarg[7]="no";
    fixarg[8]="0.";
    modify->add_fix(9,const_cast<char**>(fixarg));
    fix_source=static_cast<FixPropertyAtom*>(
        modify->find_fix_property(source_name,"property/atom","scalar",0,0,style));
  }
```

```
  updatePtrs();
}

/* ----------------------------------------------------------------------- */

double* FixScalarTransportEquation::get_capacity()
{
    return capacity;
}

/* ----------------------------------------------------------------------- */

void FixScalarTransportEquation::init()
{
  if (!atom->rmass_flag) error->all(FLERR,
    "Please use an atom style that defines per-particle mass
    for fix transportequation/scalar");

  if (strcmp(update->integrate_style,"respa") == 0)
    nlevels_respa = ((Respa *) update->integrate)->nlevels;

  if(capacity_flag)
  {
      int max_type = atom->get_properties()->max_type();

      if(capacity) delete []capacity;
      capacity = new double[max_type+1];
```

```
    fix_capacity = static_cast<FixPropertyGlobal*>(
      modify->find_fix_property(capacity_name,"property/global",
      "peratomtype",max_type,0,style));

    //pre-calculate parameters for possible contact material combinations
    for(int i=1;i< max_type+1; i++)
        capacity[i] = fix_capacity->compute_vector(i-1);
  }
}

/* ---------------------------------------------------------------------- */

int FixScalarTransportEquation::modify_param(int narg, char **arg)
{
  if (strcmp(arg[0],"integrate") == 0) {
    if (narg < 2) error->fix_error(FLERR,this,
        "not enough arguments for fix_modify 'integrate'");

    if (strcmp(arg[1],"start") == 0) {
      int_flag = true;
    } else if (strcmp(arg[1],"stop") == 0) {
      int_flag = false;
    } else
      error->fix_error(FLERR,this,"wrong argument for fix_modify 'integrate'");
    return 2;
  }
```

```
  if (strcmp(arg[0],"every") == 0) {
    if (narg < 2) error->fix_error(FLERR,this,
        "not enough arguments for fix_modify 'every'");

    nevery_ = force->inumeric(FLERR,arg[1]);

    return 1;
  }
  return 0;
}

/* ---------------------------------------------------------------------- */

// FixScalarTransportEquation executes cellular reactions and growth
// if reaction_flag is true
void FixScalarTransportEquation::initial_integrate(int vflag)
{

  updatePtrs();

  //Skip in case there was NO Integration the last time step (to keep flux in mem)
  if(!performedIntegrationLastStep_)
        return;

  // execute reaction if reaction_flag==true
  if (reaction_flag) {
```

```
//reset flux
//sources are not reset
// Initialize values
// nlocal is per processor, not equal to the total atoms in the system
int nlocal = atom->nlocal;
int *type = atom->type;
double  dt = update->dt;
double *rmass = atom->rmass;
double *radius = atom->radius;
const double three_quarters_pi = (3.0 / (4.0 * MY_PI));
const double four_thirds_pi = 4.0 * MY_PI / 3.0;
const double third = 1.0 / 3.0;
double *substrateQuantity;
// get substrate based on SubstrateName
fix_quantity_substrate = static_cast<FixPropertyAtom*>(
    modify->find_fix_property(SubstrateName,"property/atom",
    "scalar",0,0,style));
substrateQuantity = fix_quantity_substrate->vector_atom;

for (int i = 0; i < nlocal; i++)
{
       // start with 0 flux
       flux[i]=0.0;
       // check to make sure that concentrations stay positive.
       // Round to 0.0 otherwise
       if (substrateQuantity[i] < 0.0) {
```

```
 substrateQuantity[i]=0.0;
 printf("WARNING: Negative value of substrate quantity at particle '%i'.
 Rounding '%.30f' to 0.0.\n", i, substrateQuantity[i]);
}
if (quantity[i] < 0.0) {
 quantity[i]=0.0;
 printf("WARNING: Negative value of product quantity at particle '%i'.
 Rounding '%.30f' to 0.0.\n", i, quantity[i]);
}
// Check that the atom of interest belongs to the atomtype.
// If yes, perform reaction
if (atomtype == type[i])
{
     // Calculate specific growth rate and substrate utilization
     // rate using Monod equation
     double monod_rate = u_max *
         substrateQuantity[i] / (K_S + substrateQuantity[i]);
     // Calculate amount of biomass growth based on monod_rate
     double drmass = rmass[i] * monod_rate * double(nevery_) * dt;

     // Calculate rate of change of substrate based on biomass consumption
     double dsubstrate = drmass / yield_growth_coefficient;
     // Ensure we don't consume more substrate than available
     if (dsubstrate > substrateQuantity[i]) {
         dsubstrate = substrateQuantity[i];
         // update drmass if necessary to match
         // the reduction in substrate consumption
```

```
                drmass = dsubstrate * yield_growth_coefficient;
            }
            // Consume substrate
            substrateQuantity[i] -= dsubstrate;

            // Produce product based on amount of substrate consumed.
            // Source is the rate of produce produced
            source[i] = dsubstrate * yield_product_from_substrate /
                (double(nevery_) * dt);

            // Calculate initial biomass density
            double density = rmass[i] /
                (four_thirds_pi * radius[i] * radius[i] * radius[i]);
            // Grow by drmass
            rmass[i] += drmass;
            // Grow radius at constant density
            radius[i] = pow(three_quarters_pi * (rmass[i] / density), third);
        }
    }
// Otherwise, no reaction flag. Initialize flux at 0, as indicated by original code
} else {
    int nlocal = atom->nlocal;
    for (int i = 0; i < nlocal; i++)
    {
        flux[i]=0.0;
    }
}
```

```
  fix_quantity->do_forward_comm();
}

/* ----------------------------------------------------------------------- */

void FixScalarTransportEquation::pre_force(int vflag)
{

    if(neighbor->ago == 0)
        fix_quantity->do_forward_comm();
}

/* ----------------------------------------------------------------------- */

void FixScalarTransportEquation::final_integrate()
{

    // skip if integration turned off
    if(!int_flag)
        return;

    // skip if integration not wanted at this timestep
    if (update->ntimestep % nevery_)
    {
        performedIntegrationLastStep_ = false;
        return;
```

```
    }

    updatePtrs();

    fix_source->do_forward_comm();

    (this->*advanceQty)();

    performedIntegrationLastStep_ = true;
}

/* ----------------------------------------------------------------------- */
void FixScalarTransportEquation::advanceQtyExplicit()
{
    double  dt = update->dt;
    int     nlocal = atom->nlocal;
    double  *rmass = atom->rmass;
    int     *type = atom->type;
    int     *mask = atom->mask;
    double  capacity;

    if(capacity_flag)
    {
        for (int i = 0; i < nlocal; i++)
        {
           if (mask[i] & groupbit){
              capacity = fix_capacity->compute_vector(type[i]-1);
```

```
            if(fabs(capacity) > SMALL) quantity[i] += (
                  //multiply source to account for missing steps
                  flux[i] + source[i]*double(nevery_)
                                                    ) * dt
                                                  / (rmass[i]*capacity);
          }
        }
    }
    else
    {
        for (int i = 0; i < nlocal; i++)
        {
           if (mask[i] & groupbit){
              quantity[i] += (
                //multiply source to account for missing steps
                flux[i] + source[i]*double(nevery_)
                             ) * dt ;
           }
        }
    }
}

/* ---------------------------------------------------------------------- */
void FixScalarTransportEquation::advanceQtyImplicit()
{

    double  dt = update->dt;
```

```
int     nlocal = atom->nlocal;
double  *rmass = atom->rmass;
int     *type = atom->type;
int     *mask = atom->mask;
double  *radius  = atom->radius;
double  capacity;
double  OneMinusCN = 1.0 - crankNicholsonFactor_;

if(capacity_flag)
{
    for (int i = 0; i < nlocal; i++)
    {
       if (mask[i] & groupbit){
          capacity = fix_capacity->compute_vector(type[i]-1);
          if(fabs(capacity) > SMALL)
          {
                 double currRadius = radius[i];
                 double termM  = dt / (rmass[i]*capacity);
                 double termMP = termM
                              * transCoeffQty_[i]
                              * currRadius * currRadius * 12.5663706144 ;
                              //surface area
                 quantity[i]  = (   quantity[i]
                                 * (1.0 - termMP * OneMinusCN)
                                 + termMP * fluidQty_[i]
                                 + termM
                                 * (  flux[i]
```

```
                                 + source[i]*double(nevery_)
                                )
                              ) / (1.0 + termMP*crankNicholsonFactor_);
            }
        }
    }
}
else
{
    for (int i = 0; i < nlocal; i++)
    {
        if (mask[i] & groupbit)
        {
                  double currRadius = radius[i];
                  double termMP = dt
                                * transCoeffQty_[i]
                                * currRadius * currRadius * 12.5663706144 ;
                                //surface area
                  quantity[i]  = (   quantity[i]
                                   * (1.0 - termMP * OneMinusCN)
                                   + termMP * fluidQty_[i]
                                   + dt
                                   * (  flux[i]
                                      + source[i]*double(nevery_)
                                     )
                                 ) / (1.0 + termMP*crankNicholsonFactor_);
        }
```

```
        }
    }
}

/* ---------------------------------------------------------------------- */

void FixScalarTransportEquation::
    initial_integrate_respa(int vflag, int ilevel, int flag)
{
  // outermost level - update
  // all other levels - nothing

  if (ilevel == nlevels_respa-1) initial_integrate(vflag);
}

/* ---------------------------------------------------------------------- */

double FixScalarTransportEquation::compute_scalar()
{
    double *rmass = atom->rmass;
    int *type = atom->type;
    int nlocal = atom->nlocal;
    double capacity;

    updatePtrs();

    double quantity_sum = 0.;
```

```
    if(capacity_flag)
    {
        for (int i = 0; i < nlocal; i++)
        {
           capacity = fix_capacity->compute_vector(type[i]-1);
           quantity_sum += capacity * rmass[i] * quantity[i];

        }
    }
    else
    {
        for (int i = 0; i < nlocal; i++)
        {
           quantity_sum += quantity[i];
        }
    }

    MPI_Sum_Scalar(quantity_sum,world);

    return quantity_sum;
}

/* ---------------------------------------------------------------------- */

bool FixScalarTransportEquation::match_equation_id(const char* id)
{
```

```
    if(strcmp(id,equation_id)) return false;
    return true;
}
```

**fix_scalar_transport_equation.h**

```
/* ----------------------------------------------------------------------

    This is the LIGGGHTS

    DEM simulation engine, released by

    DCS Computing Gmbh, Linz, Austria

    http://www.dcs-computing.com,


    LIGGGHTS® is part of CFDEM®project:

    http://www.liggghts.com | http://www.cfdem.com


    Core developer and main author:

    Christoph Kloss,

    Modified for biological cell growth by:

    Andrew P. Latham, Emmanuel N. Skountzos, Tyler Quarton,

    and Rocky An


    LIGGGHTS® is open-source, distributed under the terms of the GNU Public

    License, version 2 or later. It is distributed in the hope that it will

    be useful, but WITHOUT ANY WARRANTY; without even the implied warranty

    of MERCHANTABILITY or FITNESS FOR A PARTICULAR PURPOSE. You should have

    received a copy of the GNU General Public License along with LIGGGHTS®.

    If not, see http://www.gnu.org/licenses . See also top-level README

    and LICENSE files.


    LIGGGHTS® and CFDEM® are registered trade marks of DCS Computing GmbH,

    the producer of the LIGGGHTS® software and the CFDEM®coupling software

    See http://www.cfdem.com/terms-trademark-policy for details.
```

```
------------------------------------------------------------------------
    Contributing author and copyright for this file:
    (if not contributing author is listed, this file has been contributed
    by the core developer)

    Copyright 2012-     DCS Computing GmbH, Linz
    Copyright 2009-2012 JKU Linz

    Implementation of implicit update algorithm
    Copyright 2016      TU Graz, Stefan Radl
    Copyright 2016      DCS Computing GmbH, Linz
------------------------------------------------------------------------- */

#ifdef FIX_CLASS

FixStyle(transportequation/scalar,FixScalarTransportEquation)

#else

#ifndef LMP_FIX_SCALAR_TRANSPORT_EQUATION_H
#define LMP_FIX_SCALAR_TRANSPORT_EQUATION_H

#include "fix.h"

namespace LAMMPS_NS {
```

```
class FixScalarTransportEquation : public Fix {
 public:
  FixScalarTransportEquation(class LAMMPS *, int, char **);
  ~FixScalarTransportEquation();

  virtual int setmask();
  virtual void post_create();
  virtual void pre_delete(bool unfixflag);
  virtual void init();
  virtual int modify_param(int narg, char **arg);
  virtual void updatePtrs();
  virtual void initial_integrate_respa(int,int,int);
  virtual void initial_integrate(int);
  virtual void pre_force(int vflag);
  virtual void final_integrate();
  virtual double compute_scalar();
  bool match_equation_id(const char*);

  //Tools for implicit fix handling
  bool isImplicit() { return implicitMode_;};
  void register_implicit_fixes(char*, double, char*, double);
  void updatePtrsImpl();

  double *get_capacity();

  inline int n_every()
  { return nevery_; }
```

```
protected:

 void (FixScalarTransportEquation::*advanceQty)();
 void advanceQtyExplicit();
 void advanceQtyImplicit();

 int nlevels_respa;

 char *equation_id;

 class FixPropertyAtom* fix_quantity;
 char *quantity_name;
 // Added class for biological cell growth
 class FixPropertyAtom* fix_quantity_substrate;



 class FixPropertyAtom* fix_flux;
 char *flux_name;
 class FixPropertyAtom* fix_source;
 char *source_name;

 //Implicit fixes, names, and pointers
 double crankNicholsonFactor_;
 bool  implicitMode_;
 class FixPropertyAtom* fix_fluidQty_;
 char *fluid_name_;
```

```
class FixPropertyAtom* fix_transCoeffQty_;
char *transCoeff_name_;
double *fluidQty_;
double *transCoeffQty_;
double fluidQty_0_;
double transCoeffQty_0_;

//storage capacity - would be thermal capacity for heat conduction
int capacity_flag;
class FixPropertyGlobal* fix_capacity;
double *capacity;
char *capacity_name;

double quantity_0;
double *quantity;
double *flux;
double *source;

// flag if integrate quantity or not
bool int_flag;
// flag if a biological cell growth reaction should occur or not
bool reaction_flag;

int  nevery_; //integrate only this many time steps (to avoid round-off issues)
bool performedIntegrationLastStep_;

// Added variables for biological cell growth
```

```
  double yield_product_from_substrate;
  double yield_growth_coefficient;
  int atomtype;
  char*   SubstrateName;
  double u_max;
  double K_S;
};

}

#endif
#endif
```

**cfdemSolverPisoSTM.C**

```
/*---------------------------------------------------------------------------*\
    CFDEMcoupling - Open Source CFD-DEM coupling

    CFDEMcoupling is part of the CFDEMproject
    www.cfdem.com
                                Christoph Goniva,
                                Copyright (C) 1991-2009 OpenCFD Ltd.
                                Copyright (C) 2012-     DCS Computing GmbH,Linz
    Modified by Andrew Latham to add gravitational forces to the fluid
-------------------------------------------------------------------------------
License
    This file is part of CFDEMcoupling.

    CFDEMcoupling is free software: you can redistribute it and/or modify it
    under the terms of the GNU General Public License as published by
    the Free Software Foundation, either version 3 of the License, or
    (at your option) any later version.

    CFDEMcoupling is distributed in the hope that it will be useful, but WITHOUT
    ANY WARRANTY; without even the implied warranty of MERCHANTABILITY or
    FITNESS FOR A PARTICULAR PURPOSE.  See the GNU General Public License
    for more details.

    You should have received a copy of the GNU General Public License
    along with CFDEMcoupling.  If not, see <http://www.gnu.org/licenses/>.
```

```
Application
    cfdemSolverPisoSTM

Description
    Transient solver for incompressible flow.
    Turbulence modelling is generic, i.e. laminar, RAS or LES may be selected.
    The code is an evolution of the solver pisoFoam in OpenFOAM(R) 1.6,
    where additional functionality for CFD-DEM coupling is added.
\*---------------------------------------------------------------------------*/

#include "fvCFD.H"
#include "singlePhaseTransportModel.H"

#include "OFversion.H"
#if defined(version30)
    #include "turbulentTransportModel.H"
    #include "pisoControl.H"
#else
    #include "turbulenceModel.H"
#endif
#if defined(versionv1606plus) || defined(version40)
    #include "fvOptions.H"
#else
    #include "fvIOoptionList.H"
#endif
#include "fixedFluxPressureFvPatchScalarField.H"
#ifdef MS
```

```
    #include "cfdemCloudMS.H"
#else
    #include "cfdemCloud.H"
#endif
#if defined(anisotropicRotation)
    #include "cfdemCloudRotation.H"
#endif
#include "implicitCouple.H"
#include "clockModel.H"
#include "smoothingModel.H"
#include "forceModel.H"

#include "scalarTransportModel.H"
// * * * * * * * * * * * * * * * * * * * * * * * * * * * * * * * * * * * * * * //

int main(int argc, char *argv[])
{
    #include "setRootCase.H"
    #include "createTime.H"
    #include "createMesh.H"
    #if defined(version30)
        pisoControl piso(mesh);
        #include "createTimeControls.H"
    #endif
    #include "createFields.H"
    #include "createFvOptions.H"
    #include "initContinuityErrs.H"
```

```
// create cfdemCloud
#include "readGravitationalAcceleration.H"
#include "checkImCoupleM.H"
#if defined(anisotropicRotation)
    cfdemCloudRotation particleCloud(mesh);
#else
    #ifdef MS
        cfdemCloudMS particleCloud(mesh);
    #else
        cfdemCloud particleCloud(mesh);
    #endif
#endif
#include "checkModelType.H"

// create a scalarTransportModel
autoPtr<scalarTransportModel> stm
(
    scalarTransportModel::New(particleCloud.couplingProperties(),particleCloud)
);

// * * * * * * * * * * * * * * * * * * * * * * * * * * * * * * * * * * * * * //
Info<< "\nStarting time loop\n" << endl;
while (runTime.loop())
{
    Info<< "Time = " << runTime.timeName() << nl << endl;
```

```
#if defined(version30)
    #include "readTimeControls.H"
    #include "CourantNo.H"
    #include "setDeltaT.H"
#else
    #include "readPISOControls.H"
    #include "CourantNo.H"
#endif

// do particle stuff
particleCloud.clockM().start(1,"Global");
particleCloud.clockM().start(2,"Coupling");
bool hasEvolved = particleCloud.evolve(voidfraction,Us,U);

if(hasEvolved)
{
    particleCloud.smoothingM().smoothenAbsolutField(
        particleCloud.forceM(0).impParticleForces());
}

Ksl = particleCloud.momCoupleM(particleCloud.registryM().getProperty(
    "implicitCouple_index")).impMomSource();
Ksl.correctBoundaryConditions();

surfaceScalarField voidfractionf = fvc::interpolate(voidfraction);
phi = voidfractionf*phiByVoidfraction;
```

```
        //Force Checks
        #include "forceCheckIm.H"

        #include "solverDebugInfo.H"
        particleCloud.clockM().stop("Coupling");

        particleCloud.clockM().start(26,"Flow");

        //Scalar transport if desired.
        //Use "none" (noTransport) if no scalar transport is desired
        stm().update();

        if(particleCloud.solveFlow())
        {
            // Pressure-velocity PISO corrector
            {
                // Momentum predictor
                fvVectorMatrix UEqn
                (
                    fvm::ddt(voidfraction,U) - fvm::Sp(fvc::ddt(voidfraction),U)
                  + fvm::div(phi,U) - fvm::Sp(fvc::div(phi),U)
//                + turbulence->divDevReff(U)
                  + particleCloud.divVoidfractionTau(U, voidfraction)
                  ==
                  - fvm::Sp(Ksl/rho,U)
                  + fvOptions(U)
                );
```

```
UEqn.relax();
fvOptions.constrain(UEqn);

#if defined(version30)
    if (piso.momentumPredictor())
#else
    if (momentumPredictor)
#endif
{
    if (modelType=="B" || modelType=="Bfull")
    // Add gravity (g) when solving the velocity
        solve(UEqn == - fvc::grad(p) + g + Ksl/rho*Us);
    else
    // Add gravity (g) when solving the velocity
        solve(UEqn == - voidfraction*fvc::grad(p) + g + Ksl/rho*Us);

    fvOptions.correct(U);
}

// --- PISO loop
#if defined(version30)
    while (piso.correct())
#else
    for (int corr=0; corr<nCorr; corr++)
#endif
{
```

```
volScalarField rUA = 1.0/UEqn.A();

surfaceScalarField rUAf("(1|A(U))", fvc::interpolate(rUA));
volScalarField rUAvoidfraction(
    "(voidfraction2|A(U))",rUA*voidfraction);
surfaceScalarField rUAfvoidfraction(
    "(voidfraction2|A(U)F)", fvc::interpolate(rUAvoidfraction));

U = rUA*UEqn.H();

#ifdef version23
    phi = ( fvc::interpolate(U) & mesh.Sf() )
        + rUAfvoidfraction*fvc::ddtCorr(U, phiByVoidfraction);
#else
    phi = ( fvc::interpolate(U) & mesh.Sf() )
        + fvc::ddtPhiCorr(rUAvoidfraction, U, phiByVoidfraction);
#endif
surfaceScalarField phiS(fvc::interpolate(Us) & mesh.Sf());
phi += rUAf*(fvc::interpolate(Ksl/rho) * phiS);

if (modelType=="A")
    rUAvoidfraction = volScalarField(
        "(voidfraction2|A(U))",rUA*voidfraction*voidfraction);

// Update the fixedFluxPressure BCs to ensure flux consistency
#include "fixedFluxPressureHandling.H"
```

```
// Non-orthogonal pressure corrector loop
#if defined(version30)
    while (piso.correctNonOrthogonal())
#else
    for (int nonOrth=0; nonOrth<=nNonOrthCorr; nonOrth++)
#endif
{
    // Pressure corrector
    fvScalarMatrix pEqn
    (
        fvm::laplacian(rUAvoidfraction, p) ==
            fvc::div(voidfractionf*phi) +
            particleCloud.ddtVoidfraction()
    );
    pEqn.setReference(pRefCell, pRefValue);

    #if defined(version30)
        pEqn.solve(mesh.solver(p.select(piso.finalInnerIter())));
        if (piso.finalNonOrthogonalIter())
        {
            phiByVoidfraction = phi - pEqn.flux()/voidfractionf;
        }
    #else
        if( corr == nCorr-1 && nonOrth == nNonOrthCorr )
            #if defined(versionExt32)
                pEqn.solve(mesh.solutionDict().solver("pFinal"));
            #else
```

```
                    pEqn.solve(mesh.solver("pFinal"));
                #endif
            else
                pEqn.solve();

            if (nonOrth == nNonOrthCorr)
            {
                phiByVoidfraction = phi - pEqn.flux()/voidfractionf;
            }
        #endif

    } // end non-orthogonal corrector loop

    phi = voidfractionf*phiByVoidfraction;
    #include "continuityErrorPhiPU.H"

    if (modelType=="B" || modelType=="Bfull")
        U -= rUA*fvc::grad(p) - Ksl/rho*Us*rUA;
    else
        U -= voidfraction*rUA*fvc::grad(p) - Ksl/rho*Us*rUA;

    U.correctBoundaryConditions();
    fvOptions.correct(U);

} // end piso loop
}
```

```
            laminarTransport.correct();
            turbulence->correct();
        }// end solveFlow
        else
        {
            Info << "skipping flow solution." << endl;
        }

        runTime.write();

        Info<< "ExecutionTime = " << runTime.elapsedCpuTime() << " s"
            << "  ClockTime = " << runTime.elapsedClockTime() << " s"
            << nl << endl;

        particleCloud.clockM().stop("Flow");
        particleCloud.clockM().stop("Global");
    }

    Info<< "End\n" << endl;

    return 0;
}


// ************************************************************************* //
```

#### Analysis and Initial Configurations

The script analyze_DEM.py was used to analyze the output of the simulations. It assumes 2 types of particles ("1" and "2") and 2 metabolites ("C" and "D"). It analyzes dump files, written as dump*.vtk, where * is the timestep. The variables min_value, max_value, and step can be changed to determine which dump files to read. The current script will analyze and write output to the current working directory. The results can be written to a different directory by changing the results_folder variable in main, while changing data_dir can tell the code to look for the dump*.vtk files in a different directory.

The code writes five output files:

1. volumes.dat - has 4 columns: i) the timestep, ii) the average volume change of all cells, iii) the average volume change of type 1 cells, and iv) the average volume change of type 2 cells. Volume changes assume the simulation starts from a sphere with a radius of 0.56.

2. radius.dat - has 3 columns: i) the timestep, ii) the average radius of type 1 cells, and iii) the average radius of type 2 cells.

3. average_metabolites.dat - has 7 columns: i) the timestep, ii) the average amount of C in all cells, iii) the average amount of D in all cells, iv) the minimum value of C in all cells, v) the minimum value of D in all cells, vi) the maximum of C in all cells, and vii) the maximum of D in all cells.

4. source_metabolites.dat - has 7 columns: i) the timestep, ii) the average amount of CSource in all cells, iii) the average amount of DSource in all cells, iv) the minimum value of CSource in all cells, v) the minimum value of DSource in all cells, vi) the maximum of CSource in all cells, and vii) the maximum of DSource in all cells.

5. flux_metabolites.dat - has 7 columns: i) the timestep, ii) the average amount of CFlux in all cells, iii) the average amount of DFlux in all cells, iv) the minimum value

of CFlux in all cells, v) the minimum value of DFlux in all cells, vi) the maximum of CFlux in all cells, and vii) the maximum of DFlux in all cells.

Code for writing initial configurations for CAMDLES2 simulations is also included. The 3 scripts (`write_2colony.py`, `write_data_sedimentation.py`, and `write_data_uniform.py`) correspond to each of the 3 starting configurations used in the paper. Each script is run in python, requires no arguments, and requires MDAnalysis (https://www.mdanalysis.org/) as a prerequisite. The script will write a liggghts data file titled `liggghts.data` in the current working directory.

**analyze_DEM.py**

```
# Script to analyze CAMDLES simulations of microbial cell growth.
# Assumes 2 types of particles ("1" and "2"), and 2 metabolites ("C" and "D")
# Analyzes dump files, written as dump*.vtk, where * is the timestep.
# Change min_value, max_value, and step to determine which dump files to read
# The current script will analyze and write output to the current working directory.
# The results can be written to a different directory by chaning the
# results_folder variable in main, while chaning data_dir
# can tell the code to look for the dump*.vtk files in a different directory
# Writes 5 output files:
# 1. volumes.dat - has 4 columns: i) the timestep, ii) the average volume
# change of all cells, iii) the average volume change of type 1 cells,
# and iv) the average volume change of type 2 cells.
# Volume changes assume the simulation starts from a sphere with a radius of 0.56.
# 2. radius.dat - has 3 columns: i) the timestep, ii) the average radius of
# type 1 cells, and iii) the average radius of type 2 cells
# 3. average_metabolites.dat - has 7 columns: i) the timestep,
# ii) the average amount of C in all cells, iii) the average amount of
# D in all cells, iv) the minimum value of C in all cells,
# v) the minimum value of D in all cells, vi) the maximum of C in all cells,
# and vii) the maximum of D in all cells.
# 4. source_metabolites.dat - has 7 columns: i) the timestep,
# ii) the average amount of CSource in all cells, iii) the average amount of
# DSource in all cells, iv) the minimum value of CSource in all cells,
# v) the minimum value of DSource in all cells, vi)
# the maximum of CSource in all cells, and vii) the maximum of DSource in all cells.
# 5. flux_metabolites.dat - has 7 columns: i) the timestep,
```

```
# ii) the average amount of CFlux in all cells, iii) the average
# amount of DFlux in all cells, iv) the minimum value of
# CFlux in all cells, v) the minimum value of DFlux in all cells,
# vi) the maximum of CFlux in all cells, and vii) the maximum of DFlux in all cells.

import os
import numpy as np

# Initial volume calculation with radius = 0.56 at t = 0
V0 = (4 / 3) * np.pi * (0.56**3)



def read_vtk_files(
    data_dir,
    min_value,
    max_value,
    step,
    number_of_cells,
    avg_volume_file,
    avg_metabolites_file,
    source_metabolites_file,
    flux_metabolites_file,
    avg_radius_file,
):
    num_lines = (
        number_of_cells + 8
    ) // 9  # Calculate the number of lines to read
```

```
with open(avg_volume_file, "w") as volume_file, open(
    avg_metabolites_file, "w"
) as metabolites_file, open(
    source_metabolites_file, "w"
) as source_metabolites, open(
    flux_metabolites_file, "w"
) as flux_metabolites, open(
    avg_radius_file, "w"
) as radius_file:
    for i in range(min_value, max_value + 1, step):
        filename = f"{data_dir}dump{i}.vtk"
        if os.path.exists(filename):
            print(f"Reading file: {filename}")
            with open(filename, "r") as file:
                lines = file.readlines()
                radius_index = None
                type_index = None
                speciesC_index = None
                speciesD_index = None
                speciesCSource_index = None
                speciesDSource_index = None
                speciesCFlux_index = None
                speciesDFlux_index = None

                # Find the indices for type, radius, f_speciesC, and f_speciesD
                for index, line in enumerate(lines):
```

```
if "type" in line:
    type_index = index + 1
if "radius" in line:
    radius_index = index + 1
if "f_speciesC" in line and speciesC_index == None:
    speciesC_index = index + 1
if "f_speciesD" in line and speciesD_index == None:
    speciesD_index = index + 1
if "f_speciesCSource" in line:
    speciesCSource_index = index + 1
if "f_speciesDSource" in line:
    speciesDSource_index = index + 1
if "f_speciesCFlux" in line:
    speciesCFlux_index = index + 1
if "f_speciesDFlux" in line:
    speciesDFlux_index = index + 1
if (
    type_index
    and radius_index
    and speciesC_index
    and speciesD_index
    and speciesCSource_index
    and speciesDSource_index
    and speciesCFlux_index
    and speciesDFlux_index
):
    break
```

```
if (
    radius_index is not None
    and type_index is not None
    and speciesC_index is not None
    and speciesD_index is not None
):
    data_points_radius = []
    data_points_speciesC = []
    data_points_speciesD = []
    data_points_speciesCSource = []
    data_points_speciesDSource = []
    data_points_speciesCFlux = []
    data_points_speciesDFlux = []
    data_points_type = []
    points_read = 0

    # Extract type data
    for line in lines[type_index : type_index + num_lines]:
        values = line.strip().split()
        for value in values:
            if points_read < number_of_cells:
                data_points_type.append(int(value))
                points_read += 1
            else:
                break
    if points_read != number_of_cells:
```

```
    print(
        f"WARNING at type in {filename}. Number of cells " \
        "expected: {number_of_cells}." \
        " Points read: {points_read}"
    )

points_read = 0  # Reset for radius data

# Extract radius data
for line in lines[
    radius_index : radius_index + num_lines
]:
    values = line.strip().split()
    for value in values:
        if points_read < number_of_cells:
            data_points_radius.append(float(value))
            points_read += 1
        else:
            break
if points_read != number_of_cells:
    print(
        f"WARNING at radius in {filename}." \
        " Number of cells expected: {number_of_cells}." \
        " Points read: {points_read}"
    )

# Extract f_speciesC data
```

```
points_read = 0
for line in lines[
    speciesC_index : speciesC_index + num_lines
]:
    values = line.strip().split()
    for value in values:
        if points_read < number_of_cells:
            data_points_speciesC.append(float(value))
            points_read += 1
        else:
            break
if points_read != number_of_cells:
    print(
        f"WARNING at f_speciesC in {filename}." \
        " Number of cells expected: {number_of_cells}." \
        " Points read: {points_read}"
    )

# Extract f_speciesD data
points_read = 0
for line in lines[
    speciesD_index : speciesD_index + num_lines
]:
    values = line.strip().split()
    for value in values:
        if points_read < number_of_cells:
            data_points_speciesD.append(float(value))
```

```
            points_read += 1
        else:
            break
if points_read != number_of_cells:
    print(
        f"WARNING at f_speciesD in {filename}." \
        " Number of cells expected: {number_of_cells}." \
        " Points read: {points_read}"
    )

# Extract f_speciesCSource data
points_read = 0
for line in lines[
    speciesCSource_index : speciesCSource_index
    + num_lines
]:
    values = line.strip().split()
    for value in values:
        if points_read < number_of_cells:
            data_points_speciesCSource.append(
                float(value)
            )
            points_read += 1
        else:
            break
if points_read != number_of_cells:
    print(
```

```
        f"WARNING at f_speciesCSource in {filename}." \
        " Number of cells expected: {number_of_cells}." \
        " Points read: {points_read}"
    )

# Extract f_speciesDSource data
points_read = 0
for line in lines[
    speciesDSource_index : speciesDSource_index
    + num_lines
]:
    values = line.strip().split()
    for value in values:
        if points_read < number_of_cells:
            data_points_speciesDSource.append(
                float(value)
            )
            points_read += 1
        else:
            break
if points_read != number_of_cells:
    print(
        f"WARNING at f_speciesDSource in {filename}." \
        " Number of cells expected: {number_of_cells}." \
        " Points read: {points_read}"
    )
```

```
# Extract f_speciesCFlux data
points_read = 0
for line in lines[
    speciesCFlux_index : speciesCFlux_index + num_lines
]:
    values = line.strip().split()
    for value in values:
        if points_read < number_of_cells:
            data_points_speciesCFlux.append(
                float(value)
            )
            points_read += 1
        else:
            break
if points_read != number_of_cells:
    print(
        f"WARNING at f_speciesCFlux in {filename}." \
        " Number of cells expected: {number_of_cells}." \
        " Points read: {points_read}"
    )

# Extract f_speciesDFlux data
points_read = 0
for line in lines[
    speciesDFlux_index : speciesDFlux_index + num_lines
]:
    values = line.strip().split()
```

```
    for value in values:
        if points_read < number_of_cells:
            data_points_speciesDFlux.append(
                float(value)
            )
            points_read += 1
        else:
            break
if points_read != number_of_cells:
    print(
        f"WARNING at f_speciesDFlux in {filename}." \
        " Number of cells expected: {number_of_cells}." \
        " Points read: {points_read}"
    )

# Loop over radius. Check for negatives
for idx, value in enumerate(
    data_points_radius, start=1
):
    if value < 0:
        print(
            f"WARNING!!! Negative radius " \
            "({value}) in {filename}."
        )

# Calculate average radius and volume for all cells
ave_radius = np.mean(data_points_radius)
```

```
ave_volume_all = (4 / 3) * np.pi * (ave_radius**3)

# Calculate average radius and volume
# for type 1 and type 2 cells
radius_type1 = [
    r
    for r, t in zip(
        data_points_radius, data_points_type
    )
    if t == 1
]
radius_type2 = [
    r
    for r, t in zip(
        data_points_radius, data_points_type
    )
    if t == 2
]

ave_radius_type1 = (
    np.mean(radius_type1) if radius_type1 else 0
)
ave_radius_type2 = (
    np.mean(radius_type2) if radius_type2 else 0
)
ave_volume_type1 = (
    (4 / 3) * np.pi * (ave_radius_type1**3)
```

```
    if ave_radius_type1
    else 0
)
ave_volume_type2 = (
    (4 / 3) * np.pi * (ave_radius_type2**3)
    if ave_radius_type2
    else 0
)

# Fractional increase in volume
fract_incr_volume_all = ave_volume_all / V0
fract_incr_volume_type1 = (
    ave_volume_type1 / V0
    if ave_volume_type1 != 0
    else 0
)
fract_incr_volume_type2 = (
    ave_volume_type2 / V0
    if ave_volume_type2 != 0
    else 0
)

# Write to the average volumes file
volume_file.write(
    f"{i} {fract_incr_volume_all} " \
        "{fract_incr_volume_type1} {fract_incr_volume_type2}\n"
)
```

```
# Write average radius to the average_radius file
radius_file.write(
    f"{i} {ave_radius_type1} {ave_radius_type2}\n"
)

# Calculate averages for f_speciesC and f_speciesD
ave_speciesC = np.mean(data_points_speciesC)
ave_speciesD = np.mean(data_points_speciesD)
min_speciesC = np.min(data_points_speciesC)
max_speciesC = np.max(data_points_speciesC)
min_speciesD = np.min(data_points_speciesD)
max_speciesD = np.max(data_points_speciesD)

# Write to the average metabolites file
metabolites_file.write(
    f"{i} {ave_speciesC} {ave_speciesD}" \
        " {min_speciesC} {min_speciesD}" \
        " {max_speciesC} {max_speciesD}\n"
)

# Calculate averages for f_speciesCSource and f_speciesDSource
ave_speciesCSource = np.mean(
    data_points_speciesCSource
)
ave_speciesDSource = np.mean(
    data_points_speciesDSource
```

```
)
min_speciesCSource = np.min(data_points_speciesCSource)
max_speciesCSource = np.max(data_points_speciesCSource)
min_speciesDSource = np.min(data_points_speciesDSource)
max_speciesDSource = np.max(data_points_speciesDSource)

# Write to the average metabolites file
source_metabolites.write(
    f"{i} {ave_speciesCSource} {ave_speciesDSource}" \
        " {min_speciesCSource} {min_speciesDSource}" \
        " {max_speciesCSource} {max_speciesDSource}\n"
)

# Calculate averages for f_speciesCFlux and f_speciesDFlux
ave_speciesCFlux = np.mean(data_points_speciesCFlux)
ave_speciesDFlux = np.mean(data_points_speciesDFlux)
min_speciesCFlux = np.min(data_points_speciesCFlux)
max_speciesCFlux = np.max(data_points_speciesCFlux)
min_speciesDFlux = np.min(data_points_speciesDFlux)
max_speciesDFlux = np.max(data_points_speciesDFlux)

# Write to the average metabolites file
flux_metabolites.write(
    f"{i} {ave_speciesCFlux} {ave_speciesDFlux}" \
        " {min_speciesCFlux} {min_speciesDFlux}" \
        " {max_speciesCFlux} {max_speciesDFlux}\n"
)
```

```
                    else:
                        print(
                            f"Required data ('type', 'radius'," \
                                " 'f_speciesC', 'f_speciesD') not found in {filename}"
                        )
            else:
                print(f"File not found: {filename}")


def main():
    try:
        results_folder = ""
        # paramters related to the simulation duration
        min_value = int(0)
        max_value = int(100000)
        step = int(1000)
        number_of_cells = int(2000)
        # pick the correct path for results
        avg_volume_file = f"{results_folder}volumes.dat"
        avg_metabolites_file = f"{results_folder}average_metabolites.dat"
        source_metabolites_file = f"{results_folder}source_metabolites.dat"
        flux_metabolites_file = f"{results_folder}flux_metabolites.dat"
        avg_radius_file = f"{results_folder}radius.dat"
        data_dir = ""
        read_vtk_files(
            data_dir,
```

```
            min_value,
            max_value,
            step,
            number_of_cells,
            avg_volume_file,
            avg_metabolites_file,
            source_metabolites_file,
            flux_metabolites_file,
            avg_radius_file,
        )
        print(
            f"Data has been written to {avg_volume_file}, " \
                "{avg_metabolites_file}, {source_metabolites_file}," \
                " {flux_metabolites_file}, {avg_radius_file}."
        )
    except ValueError:
        print(
            "Please enter valid integer values for " \
                "min, max, step, and number_of_cells."
        )


if __name__ == "__main__":
    main()
```

**write_data_uniform.py**

```
import sys
from MDAnalysis.analysis.distances import distance_array
import numpy as np
import random



# Function to write a lammps data file.
# Writes atoms uniformally distributed in a simulation box (xyz)
# natoms1 - number of atoms of type 1
# natoms2 - number of atoms of type 2
# xyz - x, y, and z dimensions of the simulation box
# R - radius of the atoms
# density - density of the atoms
# output_fn - output file name
def write_dat(
    natoms1=1333,
    natoms2=667,
    xyz=[300, 300, 300],
    R=0.56,
    density="1.08000000000000000e+00",
    output_fn="liggghts.data",
):
    # Initialize file
    new = open(output_fn, "w")
    # Write header
    new.write("LIGGGHTS data file, writen by write_data.py\n\n")
```

```
natoms = natoms1 + natoms2
new.write(f"{natoms} atoms\n")
new.write(f"2 atom types\n\n")
# Write box
new.write(f"{0:.17e} {xyz[0]:.17e} xlo xhi\n")
new.write(f"{0:.17e} {xyz[1]:.17e} ylo yhi\n")
new.write(f"{0:.17e} {xyz[2]:.17e} zlo zhi\n\n")
# Determine atom positions
pos_list = []
for i in range(natoms):
    # Add first atom
    if i == 0:
        pos = [
            random.uniform(0, xyz[0]),
            random.uniform(0, xyz[1]),
            random.uniform(0, xyz[2]),
        ]
        pos_list.append(pos)
    # For all other atoms, prevent overlap
    else:
        dist = -1
        while dist < 2 * R:
            pos = [
                random.uniform(0, xyz[0]),
                random.uniform(0, xyz[1]),
                random.uniform(0, xyz[2]),
            ]
```

```
            dist_mat = distance_array(
                np.asarray(pos), np.asarray(pos_list)
            )
            dist = np.min(dist_mat)
        # Only add postion to the list if it does not overlap with other particles
        pos_list.append(pos)
# Add type 1 atoms
new.write("Atoms\n\n")
index = 0
# Set precision of diameter manually to avoid rounding errors
D = f"{2*R:.10f}0000000e+00"
for i in range(natoms1):
    new.write(
        f"{index+1} 1 {D} {density} {pos_list[index][0]:.17e}" \
            " {pos_list[index][1]:.17e} {pos_list[index][2]:.17e} 0 0 0\n"
    )
    index += 1
# Add type 2 atoms
for i in range(natoms2):
    new.write(
        f"{index+1} 2 {D} {density} {pos_list[index][0]:.17e}" \
            " {pos_list[index][1]:.17e} {pos_list[index][2]:.17e} 0 0 0\n"
    )
    index += 1
new.close()
```

```
if __name__ == "__main__":
    write_dat()
```

**write_data_2colony.py**

```
import sys

from MDAnalysis.analysis.distances import distance_array

import numpy as np

import random



# Function to write a lammps data file.

# Writes atoms in 2 seperate boxes (xyz1 and xyz2)

# within a larger simulation box (xyz)

# natoms1 - number of atoms of type 1

# natoms2 - number of atoms of type 2

# xyz - x, y, and z dimensions of the simulation box

# xyz1 - x, y, and z dimensions of the portion of the box containing type 1 atoms

# xyz2 - x, y, and z dimensions of the portion of the box containing type 2 atoms

# R - radius of the atoms

# density - density of the atoms

# output_fn - output file name

def write_dat(

    natoms1=1333,

    natoms2=667,

    xyz=[300, 300, 300],

    xyz1=[[25, 125], [25, 125], [25, 125]],

    xyz2=[[175, 275], [175, 275], [175, 275]],

    R=0.56,

    density="1.08000000000000000e+00",

    output_fn="liggghts.data",
```

```
):
    # Initialize file
    new = open(output_fn, "w")
    # Write header
    new.write("LIGGGHTS data file, writen by write_data.py\n\n")
    natoms = natoms1 + natoms2
    new.write(f"{natoms} atoms\n")
    new.write(f"2 atom types\n\n")
    # Write box
    new.write(f"{0:.17e} {xyz[0]:.17e} xlo xhi\n")
    new.write(f"{0:.17e} {xyz[1]:.17e} ylo yhi\n")
    new.write(f"{0:.17e} {xyz[2]:.17e} zlo zhi\n\n")
    # Determine atom positions for type 1 atoms
    pos_list = []
    for i in range(natoms1):
        # Add first atom
        if i == 0:
            pos = [
                random.uniform(xyz1[0][0], xyz1[0][1]),
                random.uniform(xyz1[1][0], xyz1[1][1]),
                random.uniform(xyz1[2][0], xyz1[2][1]),
            ]
            pos_list.append(pos)
        # For all other atoms, prevent overlap
        else:
            dist = -1
            while dist < 2 * R:
```

```
            pos = [
                random.uniform(xyz1[0][0], xyz1[0][1]),
                random.uniform(xyz1[1][0], xyz1[1][1]),
                random.uniform(xyz1[2][0], xyz1[2][1]),
            ]
            dist_mat = distance_array(
                np.asarray(pos), np.asarray(pos_list)
            )
            dist = np.min(dist_mat)
        # Only add postion to the list if it does not overlap with other particles
        pos_list.append(pos)
# Determine atom positions for type 2 atoms
for i in range(natoms2):
    # Add first atom
    if i == 0:
        pos = [
            random.uniform(xyz2[0][0], xyz2[0][1]),
            random.uniform(xyz2[1][0], xyz2[1][1]),
            random.uniform(xyz2[2][0], xyz2[2][1]),
        ]
        pos_list.append(pos)
    # For all other atoms, prevent overlap
    else:
        dist = -1
        while dist < 2 * R:
            pos = [
                random.uniform(xyz2[0][0], xyz2[0][1]),
```

```
                random.uniform(xyz2[1][0], xyz2[1][1]),
                random.uniform(xyz2[2][0], xyz2[2][1]),
            ]
            dist_mat = distance_array(
                np.asarray(pos), np.asarray(pos_list)
            )
            dist = np.min(dist_mat)
        # Only add postion to the list if it does not overlap with other particles
        pos_list.append(pos)
# Add type 1 atoms
new.write("Atoms\n\n")
index = 0
# Set precision of diameter manually to avoid rounding errors
D = f"{2*R:.10f}0000000e+00"
for i in range(natoms1):
    new.write(
        f"{index+1} 1 {D} {density} {pos_list[index][0]:.17e" \
            "} {pos_list[index][1]:.17e} {pos_list[index][2]:.17e} 0 0 0\n"
    )
    index += 1
# Add type 2 atoms
for i in range(natoms2):
    new.write(
        f"{index+1} 2 {D} {density} {pos_list[index][0]:.17e}" \
            " {pos_list[index][1]:.17e} {pos_list[index][2]:.17e} 0 0 0\n"
    )
    index += 1
```

```
    new.close()



if __name__ == "__main__":
    write_dat()
```

**write_data_sedimentation.py**

```
import sys
from MDAnalysis.analysis.distances import distance_array
import numpy as np
import random



# Function to write a lammps data file. Writes atoms sedimented
# in a specific region of the simulaiton box (xyz_particles).
# natoms1 - number of atoms of type 1
# natoms2 - number of atoms of type 2
# xyz_box - x, y, and z dimensions of the simulation box
# xyz_particles - x, y, and z dimensions of where the cells will be
# R - radius of the atoms
# density - density of the atoms
# output_fn - output file name
def write_dat(
    natoms1=1333,
    natoms2=667,
    xyz_box=[300, 300, 300],
    xyz_particles=[[0, 300], [0, 300], [0, 75]],
    R=0.56,
    density="1.0800000000000000e+00",
    output_fn="liggghts.data",
):
    # Initialize file
    new = open(output_fn, "w")
```

```
# Write header
new.write("LIGGGHTS data file, writen by write_data.py\n\n")
natoms = natoms1 + natoms2
new.write(f"{natoms} atoms\n")
new.write(f"2 atom types\n\n")
# Write box
new.write(f"{0:.17e} {xyz_box[0]:.17e} xlo xhi\n")
new.write(f"{0:.17e} {xyz_box[1]:.17e} ylo yhi\n")
new.write(f"{0:.17e} {xyz_box[2]:.17e} zlo zhi\n\n")
# Determine atom positions
pos_list = []
for i in range(natoms):
    # Add first atom
    if i == 0:
        pos = [
            random.uniform(xyz_particles[0][0], xyz_particles[0][1]),
            random.uniform(xyz_particles[1][0], xyz_particles[1][1]),
            random.uniform(xyz_particles[2][0], xyz_particles[2][1]),
        ]
        pos_list.append(pos)
    # For all other atoms, prevent overlap
    else:
        dist = -1
        while dist < 2 * R:
            pos = [
                random.uniform(xyz_particles[0][0], xyz_particles[0][1]),
                random.uniform(xyz_particles[1][0], xyz_particles[1][1]),
```

```
                random.uniform(xyz_particles[2][0], xyz_particles[2][1]),
            ]
            dist_mat = distance_array(
                np.asarray(pos), np.asarray(pos_list)
            )
            dist = np.min(dist_mat)
        # Only add postion to the list if it does not overlap with other particles
        pos_list.append(pos)
# Add type 1 atoms
new.write("Atoms\n\n")
index = 0
# Set precision of diameter manually to avoid rounding errors
D = f"{2*R:.10f}0000000e+00"
for i in range(natoms1):
    new.write(
        f"{index+1} 1 {D} {density} {pos_list[index][0]:.17e}" \
            " {pos_list[index][1]:.17e} {pos_list[index][2]:.17e} 0 0 0\n"
    )
    index += 1
# Add type 2 atoms
for i in range(natoms2):
    new.write(
        f"{index+1} 2 {D} {density} {pos_list[index][0]:.17e}" \
            " {pos_list[index][1]:.17e} {pos_list[index][2]:.17e} 0 0 0\n"
    )
    index += 1
new.close()
```

```
if __name__ == "__main__":
    write_dat()
```

#### Example

Below, we list the input files needed for running simulations of microbial growth with CAMDLES2. The example below is for a system with no gravity, low starting metabolite concentration, and periodic boundary conditions.

Simulations are run by executing `source Allrun.sh > display`, which will run `Allrun.sh`, `parCFDDEMrun.sh`, and `parDEMrun.sh`. These scripts will use inputs divided between two directories, `CFD` for input parameters pertaining to CFD and CFD-DEM coupling and `DEM` for input parameters pertaining to DEM. Simulations will produce `dump*.vtk`, where * is the timestep. Each file represents the system at that timestep and includes the cell position, radius, and metabolite concentrations inside the cell. Simulations will also produce a series of numbered directories in the CFD folder. Each directory is titled according to the simulation time and includes files describing the state of each field. Both the dump*.vtk files and the CFD output can be visualized in Paraview (https://www.paraview.org/).

Gravitational effects, starting metabolite concentration, and periodic boundary conditions are adjustable. Gravity is changed by changing `value` in `CFD/constant/g`. Starting metabolite concentration is changed by modifying `internalField` in `CFD/0/C` and `CFD/0/D` and `species0` in `DEM/in.liggghts_resume`. Periodic boundary conditions are changed by modifying `boundary` (to `f f f`) in `DEM/in.liggghts_resume` and `DEM/in.liggghts_init`, `type` of `boundary` (to `wall`) in `CFD/system/blockMeshDict`, and `type` of `boundaryField` (to `zeroGradient`) for each field in `CFD/0`.

**Allrun.sh**

```
#!/bin/bash


#- define variables
casePath="$(dirname "$(readlink -f ${BASH_SOURCE[0]})")"


# check if mesh was built
if [ -f "$casePath/CFD/constant/polyMesh/points" ]; then
    echo "mesh was built before - using old mesh"
else
    echo "mesh needs to be built"
    cd $casePath/CFD
    blockMesh
fi


. $casePath/parDEMrun.sh


#- run parallel CFD-DEM in new terminal
. $casePath/parCFDDEMrun.sh
```

**parCFDDEMrun.sh**

```
#!/bin/bash


#- source CFDEM env vars
source /swbuild/alatham/Programs/CFDEM/source_me.bash


#- include functions
source $CFDEM_SRC_DIR/lagrangian/cfdemParticle/etc/functions.sh


#------------------------------------------------------------------------------#
#- define variables
casePath="$(dirname "$(readlink -f ${BASH_SOURCE[0]})")"
logpath=$casePath
headerText="run_parallel_cfdemSolverPiso_periodicChannel_CFDDEM"
logfileName="log_$headerText"
solverName="cfdemSolverPisoSTM"
nrProcs="12"
machineFileName="none"   # yourMachinefileName | none
debugMode="off"          # on | off| strict
testHarnessPath="$CFDEM_TEST_HARNESS_PATH"
runOctave="false"
postproc="true"
cleanCase="false"
#------------------------------------------------------------------------------#


#- call function to run a parallel CFD-DEM case
parCFDDEMrun $logpath $logfileName $casePath $headerText $solverName $nrProcs \
```

```
    $machineFileName $debugMode



if [ $runOctave == "true" ]
    then
        #-------------------------------#
        #  octave

        #- change path
        cd octave

        #- rmove old graph
        rm *.png

        #- run octave
        octave --no-gui checkVolFlow.m

        #- show plot
        eog volflow.png

        #- copy log file to test harness
        cp ../../$logfileName $testHarnessPath
        cp *.png $testHarnessPath
fi

if [ $postproc == "true" ]
  then
```

```
#- keep terminal open (if started in new terminal)
echo "simulation finished? ...press enter to proceed"
#read

#- get VTK data from liggghts dump file
# cd $casePath/DEM/post
# python -i $CFDEM_LPP_DIR/lpp.py dump*.liggghts_restart

#- get VTK data from CFD sim
cd $casePath/CFD
reconstructPar -noLagrangian
#- serial run of foamToVTK
foamToVTK
#- include functions
#source $CFDEM_SRC_DIR/lagrangian/cfdemParticle/etc/functions.sh
#- pseudo parallel run of foamToVTK
#pseudoParallelRun "foamToVTK" $nrPostProcProcessors

#- start paraview
paraview

#- keep terminal open (if started in new terminal)
echo "...press enter to clean up case"
echo "press Ctr+C to keep data"
read
```

```
fi


#- clean up case
if [ $cleanCase == "true" ]
  then
    #- clean up case
    keepDEMrestart="false"
    cleanCFDEMcase $casePath/CFD $keepDEMrestart
fi
```

**parDEMrun.sh**

```
#!/bin/bash


#- source CFDEM env vars
source /swbuild/alatham/Programs/CFDEM/source_me.bash


#- include functions
source $CFDEM_SRC_DIR/lagrangian/cfdemParticle/etc/functions.sh


echo "starting DEM run in parallel..."
#--------------------------------------------------------------------------------#
#- define variables
casePath="$(dirname "$(readlink -f ${BASH_SOURCE[0]})")"
logpath="$casePath"
headerText="run_liggghts_init_DEM"
logfileName="log_$headerText"
solverName="in.liggghts_init"
nrProcs=12
machineFileName="none"   # yourMachinefileName | none
debugMode="off"
#--------------------------------------------------------------------------------#


#- call function to run DEM case
parDEMrun $logpath $logfileName $casePath $headerText $solverName $nrProcs \
    $machineFileName $debugMode
```

**CFD/0/C**

```
FoamFile
{
    version     2.0;
    format      ascii;
    class       volScalarField;
    object      C;
}
// * * * * * * * * * * * * * * * * * * * * * * * * * * * * * * * * * * * * * * //

dimensions      [1 -3 0 0 0 0 0];

internalField   uniform 2e-06;

boundaryField
{
    bottom
    {
        type            cyclic;
    }
    top
    {
        type            cyclic;
    }
    left
    {
        type            cyclic;
```

```
    }
    right
    {
        type            cyclic;
    }
    front
    {
        type            cyclic;
    }
    back
    {
        type            cyclic;
    }
}

// ************************************************************************* //
```

**CFD/0/CSource**

```
FoamFile
{
    version     2.0;
    format      ascii;
    class       volScalarField;
    object      Csource;
}
// * * * * * * * * * * * * * * * * * * * * * * * * * * * * * * * * * * * * * //

dimensions      [1 -3 -1 0 0 0 0];

internalField   uniform 0;

boundaryField
{
    bottom
    {
        type            cyclic;
    }
    top
    {
        type            cyclic;
    }
    left
    {
        type            cyclic;
```

```
    }
    right
    {
        type            cyclic;
    }
    front
    {
        type            cyclic;
    }
    back
    {
        type            cyclic;
    }
}


// ************************************************************************* //
```

**CFD/0/D**

```
FoamFile
{
    version     2.0;
    format      ascii;
    class       volScalarField;
    object      D;
}
// * * * * * * * * * * * * * * * * * * * * * * * * * * * * * * * * * * * * * * //

dimensions      [1 -3 0 0 0 0 0];

internalField   uniform 2e-06;

boundaryField
{
    bottom
    {
        type            cyclic;
    }
    top
    {
        type            cyclic;
    }
    left
    {
        type            cyclic;
```

```
    }
    right
    {
        type            cyclic;
    }
    front
    {
        type            cyclic;
    }
    back
    {
        type            cyclic;
    }
}

// ************************************************************************* //
```

**CFD/0/DSource**

```
FoamFile
{
    version     2.0;
    format      ascii;
    class       volScalarField;
    object      Dsource;
}
// * * * * * * * * * * * * * * * * * * * * * * * * * * * * * * * * * * * * * //

dimensions      [1 -3 -1 0 0 0 0];

internalField   uniform 0;

boundaryField
{
    bottom
    {
        type            cyclic;
    }
    top
    {
        type            cyclic;
    }
    left
    {
        type            cyclic;
```

```
    }
    right
    {
        type            cyclic;
    }
    front
    {
        type            cyclic;
    }
    back
    {
        type            cyclic;
    }
}


// ************************************************************************* //
```

**CFD/0/Ksl**

```
FoamFile
{
    version     2.0;
    format      ascii;
    class       volScalarField;
    object      Ksl;
}
// * * * * * * * * * * * * * * * * * * * * * * * * * * * * * * * * * * * * * * //

dimensions      [1 -3 -1 0 0 0 0];

internalField   uniform 0;

boundaryField
{
    bottom
    {
        type            cyclic;
    }
    top
    {
        type            cyclic;
    }
    left
    {
        type            cyclic;
```

```
    }
    right
    {
        type            cyclic;
    }
    front
    {
        type            cyclic;
    }
    back
    {
        type            cyclic;
    }
}


// ************************************************************************* //
```

**CFD/0/p**

```
FoamFile
{
    version     2.0;
    format      ascii;
    class       volScalarField;
    object      p;
}
// * * * * * * * * * * * * * * * * * * * * * * * * * * * * * * * * * * * * * * //

dimensions      [0 2 -2 0 0 0 0];

internalField   uniform 0.;

boundaryField
{
    bottom
    {
        type            cyclic;
    }
    top
    {
        type            cyclic;
    }
    left
    {
        type            cyclic;
```

```
    }
    right
    {
        type            cyclic;
    }
    front
    {
        type            cyclic;
    }
    back
    {
        type            cyclic;
    }
}

// ************************************************************************* //
```

**CFD/0/rho**

```
FoamFile
{
    version     2.0;
    format      ascii;
    class       volScalarField;
    object      rho;
}
// * * * * * * * * * * * * * * * * * * * * * * * * * * * * * * * * * * * * * * //

dimensions      [1 -3 0 0 0 0 0];

internalField   uniform 1.00;

boundaryField
{
    bottom
    {
        type            cyclic;
    }
    top
    {
        type            cyclic;
    }
    left
    {
        type            cyclic;
```

```
    }
    right
    {
        type            cyclic;
    }
    front
    {
        type            cyclic;
    }
    back
    {
        type            cyclic;
    }
}
// ************************************************************************* //
```

**CFD/0/U**

```
FoamFile
{
    version     2.0;
    format      ascii;
    class       volVectorField;
    object      U;
}
// * * * * * * * * * * * * * * * * * * * * * * * * * * * * * * * * * * * * * * //

dimensions      [0 1 -1 0 0 0 0];

internalField   uniform (0 0 0);

boundaryField
{
    bottom
    {
        type            cyclic;
    }
    top
    {
        type            cyclic;
    }
    left
    {
        type            cyclic;
```

```
    }
    right
    {
        type            cyclic;
    }
    front
    {
        type            cyclic;
    }
    back
    {
        type            cyclic;
    }
}

// ************************************************************************* //
```

**CFD/0/Us**

```
FoamFile
{
    version     2.0;
    format      ascii;
    class       volVectorField;
    object      Us;
}
// * * * * * * * * * * * * * * * * * * * * * * * * * * * * * * * * * * * * * * //

dimensions      [0 1 -1 0 0 0 0];

internalField   uniform (0 0 0);

boundaryField
{
    bottom
    {
        type            cyclic;
    }
    top
    {
        type            cyclic;
    }
    left
    {
        type            cyclic;
```

```
    }
    right
    {
        type            cyclic;
    }
    front
    {
        type            cyclic;
    }
    back
    {
        type            cyclic;
    }
}


// ************************************************************************* //
```

**CFD/0/voidfraction**

```
FoamFile
{
    version     2.0;
    format      ascii;
    class       volScalarField;
    object      voidfraction;
}
// * * * * * * * * * * * * * * * * * * * * * * * * * * * * * * * * * * * * * //

dimensions      [0 0 0 0 0 0 0];

internalField   uniform 1;

boundaryField
{
    bottom
    {
        type            cyclic;
    }
    top
    {
        type            cyclic;
    }
    left
    {
        type            cyclic;
```

```
    }
    right
    {
        type            cyclic;
    }
    front
    {
        type            cyclic;
    }
    back
    {
        type            cyclic;
    }
}


// ************************************************************************* //
```

**CFD/constant/couplingProperties**

```
FoamFile
{
    version         2.0;
    format          ascii;

    root            "";
    case            "";
    instance        "";
    local           "";

    class           dictionary;
    object          couplingProperties;
}

// * * * * * * * * * * * * * * * * * * * * * * * * * * * * * * * * * * * * //

//===========================================================================//
// sub-models & settings

modelType "A";

couplingInterval 1;

voidFractionModel divided;

locateModel engine;
```

```
meshMotionModel noMeshMotion;

IOModel off;

probeModel off;

dataExchangeModel twoWayMPI;

averagingModel dense;//dilute;//

clockModel off;//standardClock;//

smoothingModel off;// localPSizeDiffSmoothing;// constDiffSmoothing; //

forceModels
(
    gradPForce
    scalarGeneralExchange
    viscForce
    noDrag
);

momCoupleModels
(
    implicitCouple
);
```

```
turbulenceModelType turbulenceProperties;

//===========================================================================//
// sub-model properties

scalarGeneralExchangeProps
{
    velFieldName            "U";
    voidfractionFieldName   "voidfraction";
    tempFieldName           "T";
    partTempName            "Temp";
    // explicit coupling
    partHeatFluxName        "convectiveHeatFlux";
    // implicit coupling
    // partHeatTransCoeffName  "heatTransCoeff";
    // partHeatFluidName       "heatFluid";
    // fluid thermal conductivity of water in (pg*microm)/(micros^3*K)
    lambda                  608.9;
    // Prandtl number for water at 18 C dimensionless
    Prandtl                 7.56;
    //useLiMason              true;

    //Lists with information for each species FOR THE PARTICLES
    //MUST be in the same order as eulerian species in 'scalarTransportProperties'
    //MUST correspond to property/atom in ligghts
    // (use 'couple/cfd/speciesConvection' to auto-generate individual fields)
```

```
    partSpeciesNames
    (
        speciesC
        speciesD
    );
    partSpeciesFluxNames
    (
        speciesCFlux
        speciesDFlux
    );
    partSpeciesTransCoeffNames
    (
        none //speciesCTransCoeff
        none //speciesDTransCoeff
    );
    partSpeciesFluidNames
    (
        none //speciesCFluid
        none //speciesDFluid
    );
    DMolecular
    (
        1e-3
        1e-3
    );
}
```

```
implicitCoupleProps
{
    velFieldName "U";
    granVelFieldName "Us";
    voidfractionFieldName "voidfraction";
}

gradPForceProps
{
    pFieldName "p";
    voidfractionFieldName "voidfraction";
    velocityFieldName "U";
    interpolation true;
}

viscForceProps
{
    velocityFieldName "U";
    interpolation true;
}

noDragProps
{
    noDEMForce true;
    keepCFDForce true;
}
```

```
engineProps
{
    treeSearch true;
}

dividedProps
{
    alphaMin 0.1;
}

twoWayMPIProps
{
    liggghtsPath "../DEM/in.liggghts_resume";
}
// ************************************************************************* //
```

**CFD/constant/g**

```
FoamFile
{
    version     2.0;
    format      ascii;
    class       uniformDimensionedVectorField;
    location    "constant";
    object      g;
}
// * * * * * * * * * * * * * * * * * * * * * * * * * * * * * * * * * * * * * * //

dimensions      [0 1 -2 0 0 0 0];
// Microgravity
value           ( 0 0 0 );
// Gravity
// value           ( 0 0 -9.80665e-06 );



// ************************************************************************** //
```

**CFD/constant/liggghtsCommands**

```
FoamFile
{
    version         2.0;
    format          ascii;

    root            "";
    case            "";
    instance        "";
    local           "";

    class           dictionary;
    object          liggghtsCommands;
}

// * * * * * * * * * * * * * * * * * * * * * * * * * * * * * * * * * * * * //

liggghtsCommandModels
(
    runLiggghts
);

// ************************************************************************* //
```

**CFD/constant/scalarTransportProperties**

```
FoamFile
{
    version         2.0;
    format          ascii;

    root            "";
    case            "";
    instance        "";
    local           "";

    class           dictionary;
    object          scalarTransportProperties;
}

// * * * * * * * * * * * * * * * * * * * * * * * * * * * * * * * * * * * * * //

//===========================================================================//
// sub-models & settings

scalarTransportModel generalManual;

generalManualProps
{
    // This is the volumetric heat capacity of water in [pJ / (K microm^3)]
    cpVolumetric 4180;
    eulerianFields
```

```
    (
        C   //any concentration fields MUST be first to have correct numbering
        D
    );

    fvOptionsC {};
    fvOptionsD {};
};

// Required, but leave blank
scalarTransportModelProps
{
    eulerianFields
    (
    );
};
// ************************************************************************* //
```

**CFD/constant/transportProperties**

```
FoamFile
{
    version     2.0;
    format      ascii;
    class       dictionary;
    location    "constant";
    object      transportProperties;
}
// * * * * * * * * * * * * * * * * * * * * * * * * * * * * * * * * * * * * * //

transportModel  Newtonian;

nu              nu [ 0 2 -1 0 0 0 0 ] 1.0;

// ************************************************************************* //
```

**CFD/constant/turbulenceProperties**

```
FoamFile
{
    version     2.0;
    format      ascii;
    class       dictionary;
    location    "constant";
    object      turbulenceProperties;
}
// * * * * * * * * * * * * * * * * * * * * * * * * * * * * * * * * * * * * * //

simulationType  laminar;

// ************************************************************************* //
```

**CFD/system/blockMeshDict**

```
FoamFile
{
    version     2.0;
    format      ascii;
    class       dictionary;
    object      blockMeshDict;
}
// * * * * * * * * * * * * * * * * * * * * * * * * * * * * * * * * * * * * * * //

convertToMeters 1.;

vertices
(
    (0. 0 0)
    (300 0 0)
    (300 300 0)
    (0. 300 0)
    (0. 0 300)
    (300 0 300)
    (300 300 300)
    (0. 300 300)
);

blocks
(
    hex (0 1 2 3 4 5 6 7) (60 60 60) simpleGrading (1 1 1)
```

```
);

boundary
(
    bottom
    {
        type cyclic;
        neighbourPatch    top;
        faces
        (
            (0 1 2 3)
        );
    }

    top
    {
        type cyclic;
        neighbourPatch    bottom;
        faces
        (
            (4 5 6 7)
        );
    }

    left
    {
        type cyclic;
```

```
    neighbourPatch    right;
    faces
    (
        (0 4 7 3)
    );
}

right
{
    type cyclic;
    neighbourPatch    left;
    faces
    (
        (1 5 6 2)
    );
}

front
{
    type cyclic;
    neighbourPatch    back;
    faces
    (
        (0 1 5 4)
    );
}
```

```
    back
    {
        type cyclic;
        neighbourPatch    front;
        faces
        (
            (3 2 6 7)
        );
    }
);


}
);

// ************************************************************************* //
```

**CFD/system/controlDict**

```
FoamFile
{
    version     2.0;
    format      ascii;
    class       dictionary;
    location    "system";
    object      controlDict;
}
// * * * * * * * * * * * * * * * * * * * * * * * * * * * * * * * * * * * * * * //

application     cfdemSolverPisoSTM;

startFrom       startTime;

startTime       0;

stopAt          endTime;

endTime         10000;

deltaT          0.1;

writeControl    timeStep;

writeInterval   1000;
```

```
purgeWrite      0;

writeFormat     ascii;

writePrecision  6;

writeCompression uncompressed;

timeFormat      general;

timePrecision   6;

DimensionedConstants
{
    unitSet             micro;
    microCoeffs
    {
    universal
        {
        c c [ 0 1 -1 0 0 0 0 ] 2.99792e+8; // speed of lsight in vacuum (Âµm/Âµs)
        G G [ -1 3 -2 0 0 0 0 ] 6.67429e-20;//gravitational constant (Âµm^3/(pgÂµs^2))
        h h [ 1 2 -1 0 0 0 0 ] 6.62607e-13; // Planck's constant (pgÂµm^2/Âµs)
        }
        electromagnetic
        {
        e e [ 0 0 1 0 0 1 0 ] 4.803204e-4; // elementary charge (pC)
        }
```

```
        atomic
        {
        me me [ 1 0 0 0 0 0 0 ] 9.10938e-16; // electron mass (pg)
        mp mp [ 1 0 0 0 0 0 0 ] 1.67262e-12; // proton mass (pg)
        }
        physicoChemical
        {
        mu mu [ 1 0 0 0 0 0 0 ] 1.66054e-12; // atomic mass unit (pg)
        k k [ 1 2 -2 -1 0 0 0 ] 1.38065e-8; // Boltzman constant (pgÂµm^2/Âµs^2K)
        }
        standard
        {
        //- Standard pressure [pg/ÂµmÂµs^2 ]
        Pstd Pstd [ 1 -1 -2 0 0 0 0 ] 1e2; // 1 pg/ÂµmÂµs^2
        //- Standard temperature [degK]
        Tstd Tstd [ 0 0 0 1 0 0 0 ] 300; // should be same as in SI unit system
        }
    }
}

// ************************************************************************ //
```

**CFD/system/decomposeParDict**

```
FoamFile
{
    version     2.0;
    format      ascii;
    class       dictionary;
    location    "system";
    object      decomposeParDict;
}
// * * * * * * * * * * * * * * * * * * * * * * * * * * * * * * * * * * * * * //

numberOfSubdomains 12;

method          simple;

simpleCoeffs
{
    n               ( 4 1 3 );
    delta           0.001;
}

// ************************************************************************* //
```

**CFD/system/fvSchemes**

```
FoamFile
{
    version     2.0;
    format      ascii;
    class       dictionary;
    location    "system";
    object      fvSchemes;
}
// * * * * * * * * * * * * * * * * * * * * * * * * * * * * * * * * * * * * * * //

ddtSchemes
{
    default         backward;//Euler;
}

gradSchemes
{
    default         Gauss linear;
}

divSchemes
{
    default         Gauss linear;
}

laplacianSchemes
```

```
{
    default         Gauss linear corrected;
}

interpolationSchemes
{
    default         linear;
}

snGradSchemes
{
    default         corrected;
}

fluxRequired
{
    default         no;
    p               ;
}


// ************************************************************************* //
```

**CFD/system/fvSolution**

```
FoamFile
{
    version     2.0;
    format      ascii;
    class       dictionary;
    location    "system";
    object      fvSolution;
}
// * * * * * * * * * * * * * * * * * * * * * * * * * * * * * * * * * * * * * * //

solvers
{
    p
    {
        solver          PCG;
        preconditioner  DIC;
        tolerance       1e-06;
        relTol          0;
    }

    pFinal
    {
        solver          PCG;
        preconditioner  DIC;
        tolerance       1e-06;
        relTol          0;
```

```
    }

    U
    {
        solver          PBiCG;
        preconditioner  DILU;
        tolerance       1e-05;
        relTol          0;
    }

    UFinal
    {
        $U
    }

    "(C|D)"
    {
        solver          PBiCG;
        preconditioner  DILU;
        tolerance       1e-06;
        relTol          0;
    }
}

relaxationFactors
{
    fields
```

```
    {
     p 1.0;
    }
    equations
    {
        "U.*"           1.0;
        "k.*"           1.;
        "epsilon.*"    1.;
    }
}

PISO
{
    nCorrectors     3;
    nNonOrthogonalCorrectors 0;
    pRefCell        0;
    pRefValue       0;
}


// ************************************************************************* //
```

**DEM/in.liggghts_init**

```
echo            both
log             ../DEM/log.liggghts
thermo_log      ../DEM/post/thermo.txt

atom_style      granular
atom_modify     map array
communicate     single vel yes

variable            couple_step index 1
variable            write_traject index 1000
variable            time_step index 0.1

boundary        p p p
newton          off

units           micro
processors     4 1 3

#read the data file
read_data  ../DEM/liggghts.data
# set initial velocities
velocity all set 0.0 0.0 0.0

neighbor        4 bin
neigh_modify    delay 0 one 2000
```

```
# Material properties required for granular pair styles

fix             m1 all property/global youngsModulus peratomtype 2060 2060
fix             m2 all property/global poissonsRatio peratomtype 0.3 0.3
fix             m3 all property/global coefficientRestitution peratomtypepair &
    2 0.5 0.5 0.5 0.5
fix   m4 all property/global coefficientFriction peratomtypepair 2 0 0 0 0
soft_particles yes

# pair style
pair_style  gran model hertz tangential history # Hertzian without cohesion
pair_coeff  * *

# timestep, gravity
timestep ${time_step}

## heat transfer
fix         ftco all property/global thermalConductivity peratomtype &
    600 600. # lambda in [W/(K*m)] #except in micro
# cp in [J/(kg*K)] #exce[t in micro]
fix         ftca all property/global thermalCapacity peratomtype 4184 4184
fix         heattransfer all heat/gran initial_temperature 300

# apply nve integration to all particles that are inserted as single particles
fix     integr all nve/sphere

# screen output
```

```
compute         1 all erotate/sphere
thermo_style    custom step atoms ke c_1 vol
thermo          ${write_traject}
thermo_modify   lost error norm no
compute_modify  thermo_temp dynamic yes

#insert the first particles so that dump is not empty
run  0
dump            dmp all custom/vtk ${write_traject} ../DEM/post/dump_init*.vtk &
    id type type x y z radius
```

**DEM/in.liggghts_resume**

```
echo            both
log             ../DEM/log.liggghts
thermo_log      ../DEM/post/thermo.txt

atom_style     granular
atom_modify     map array
communicate     single vel yes

variable            couple_step index 1
variable            write_traject index 1000
variable            time_step index 0.1

boundary     p p p
newton      off

units      micro
processors     4 1 3

#read the data file
read_data  ../DEM/liggghts.data

neighbor        4 bin
neigh_modify    delay 0 one 2000

#Material properties required for new pair styles
```

```
fix             m1 all property/global youngsModulus peratomtype 2060 2060
fix             m2 all property/global poissonsRatio peratomtype 0.3 0.3
fix             m3 all property/global coefficientRestitution peratomtypepair &
    2 0.5 0.5 0.5 0.5
fix   m4 all property/global coefficientFriction peratomtypepair 2 0 0 0 0
soft_particles yes

#pair style
pair_style gran model hertz tangential history #Hertzian without cohesion
pair_coeff * *

#timestep, gravity
timestep ${time_step}

#cfd coupling
fix  cfd all couple/cfd couple_every ${couple_step} mpi
fix  cfd2 all couple/cfd/force/implicit

## heat transfer
fix         ftco all property/global thermalConductivity peratomtype &
    600 600. # lambda in [W/(K*m)] #except in micro
# cp in [J/(kg*K)] #exce[t in micro]
fix         ftca all property/global thermalCapacity peratomtype 4184 4184
fix         cfd3 all couple/cfd/convection T0 300
fix         heattransfer all heat/gran initial_temperature 300

# species transfer - type 1 consumes speciesD and produces speciesC
```

```
fix         cfd4    all couple/cfd/speciesConvection speciesName speciesC &
    species0 0.0 yield_product_from_substrate 0.5 atomtype 1 %
    SubstrateName speciesD u_max 0.0000000606 K_S 0.000002 yield_growth_coefficient 0.5

# species transfer 2 - type 2 consumes speciesC and produces speciesD
fix         cfd5    all couple/cfd/speciesConvection speciesName speciesD &
    species0 0.0 yield_product_from_substrate 0.5 atomtype 2 &
    SubstrateName speciesC u_max 0.0000000303 K_S 0.000001 yield_growth_coefficient 0.5

# Set initial concentrations
run      0

# set particle temperature for the bed
region      total block INF INF INF INF INF INF units box
set         region total property/atom Temp 300

# apply nve integration to all particles that are inserted as single particles
fix     integr all nve/sphere

#center of mass.c
compute         centerOfMass all com

#compute total dragforce
compute  dragtotal all reduce sum f_dragforce[1] f_dragforce[2] f_dragforce[3]

# sum of explicit and implicit drag force given from CFD to DEM
variable    totalDragX equal f_cfd2[1]
```

```
variable    totalDragY equal f_cfd2[2]
variable    totalDragZ equal f_cfd2[3]

# explicit drag force given from CFD to DEM
variable    explicitDragX equal c_explDrag[1]
variable    explicitDragY equal c_explDrag[2]
variable    explicitDragZ equal c_explDrag[3]

#screen output
compute  1 all erotate/sphere
thermo_style    custom step atoms ke c_1 c_centerOfMass[3] &
    c_dragtotal[1] c_dragtotal[2] c_dragtotal[3] f_cfd2[1] f_cfd2[2] f_cfd2[3]
thermo  ${write_traject}
thermo_modify lost ignore norm no
compute_modify thermo_temp dynamic yes

#insert the first particles so that dump is not empty

dump            dmp all custom/vtk ${write_traject} ../DEM/post/dump*.vtk &
    id type type x y z radius f_speciesC f_speciesD &
    f_speciesCSource f_speciesDSource f_speciesCFlux f_speciesDFlux

dump            1 all custom ${write_traject} ../DEM/post/test.dump &
    id type x y z ix iy iz radius vx vy vz
dump_modify     1 sort id append yes

run      0
```
